# Long range electromagnetic effects drive protein-protein approaches: an egg of Coulomb

**DOI:** 10.1101/2022.03.16.484570

**Authors:** Neri Niccolai, Edoardo Morandi, Davide Alocci, Alberto Toccafondi, Andrea Bernini

**Author notes:** Corresponding Author: Prof. Neri Niccolai, Department of Biotechnology, Chemistry and Pharmacy, University of Siena, Via Moro 2, I-53100 Siena, Italy.

## Abstract

Living systems cannot rely on random intermolecular approaches inside cell crowding and hidden mechanisms must be present to favor only those molecular interactions which are specifically required by biological functions. Electromagnetic messaging among proteins is here hypothesized upon the observation that charged amino acids are most commonly located in adjacent sequence positions and/or in spatial close proximity. Molecular Dynamics simulations have been used to explore possible effects arising from concerted motions of charged amino acid side chains in two protein model systems. Protein electrodynamics seems to emerge as the framework for understanding long distance protein-ligand interactions.

**Highlights:**

1. Protein surfaces are often occupied by nearby side chains bearing opposite charges;
2. Coulomb interactions determine hindered reorientations of charged side chains;
3. MD simulations suggest time scales and extents of surface charge interactions;
4. concerted motions of electric charges can yield electromagnetic protein signaling.

## 1. Introduction

In the molecular crowding typical of biological systems, protein-protein interactions cannot be driven by fortuitous encounters. Even though this statement is rather obvious, limited advances have been made to understand the mechanisms controlling long range molecular dialogs among biomolecules. In this respect, Brownian dynamics simulations can offer a big deal of information on short distance biomolecular interactions by taking into account electrostatic, hydrodynamic and hydrophobic effects [1, 2]. Afterwards, transient encounter complexes, proposed as preliminary steps for the formation of protein-protein complexes [3], can be considered for further *in silico* screenings. However, how molecular partners are driven in close proximity for establishing transient encounters and, eventually, for their complex formation remains largely unknown.

Some clues for understanding this fundamental aspect of Life at the atomic level arise from a recent investigation of our group [4]. There, protein structural layers were defined together with the corresponding amino acid compositions for a large subset of PDB structures fulfilling the following requirements: i) single chain proteins, ii) proteins exhibiting only monomer biological assembly, iii) single entity structure, that is protein structures not including any type of bound ligands, iv) X ray structures ensuring single model structures, not incorporating extensive floppy regions and v) presence at 95% identity cutoff, identifying a collection of 2,410 “protein singles”. It follows that protein structures of this PDB selection are characterized by outer protein layers which are not perturbed by intermolecular interactions different from the ones due to crystal packing. Then, by using SADIC algorithm [5], a complete atom depth analysis for all these “protein singles” has been carried out. Among the proposed onion-like structural layers, all protruding moieties of these proteins could be unambiguously identified as elements of the outer structural layer [4]. Quantitative systematic analysis on amino acid composition of the most external layer revealed that lysines and glutamates resulted to be, by far, the most abundant amino acids, with 17% and 15% occurrences respectively. This finding is not surprising, considering the role of charged surface moieties to ensure suitable solubility, structural stability and intermolecular interactions. However, the fact that the latter two amino acids were found most frequently as sequence neighbors was much less expected [4]. Furthermore, DSSP assignments [6] of EK and KE protruding fragments of protein singles clearly indicated their location in helices and, to a lesser extent, in turns. This finding suggested multiple formation of salt bridges on protein surfaces, as helices and turns allow close approaches between side chains of adjacent residues. KK and EE dipeptides were also frequently found in protein outer layer, (see Table SIII of ref. 2), sharing also the same secondary structure elements of EK and KE.

It must be underlined that only the above described Bioinformatics approach could unveil such a peculiar distribution of electric charges, leaving Computational Biology to find other clues for solving the riddle underlying intermolecular chats. Indeed, once high throughput screening of amino acid composition of protein outer structural layers has been completed, the overall electric charge network needs detailed analyses. Due to the expected large flexibility of surface exposed amino acid side chains, electric charge distribution cannot be analyzed simply by inspection of static structures. Hence, MD simulations, offering a reliable reference for investigating time dependent structural features in proteins [7], are proposed as the computational basis for characterizing the evolution of electric charge networks.

Here, the hypothesis that protein surface electrodynamics is the actual engine driving medium to long range intermolecular interactions is explored by analyzing distribution and motion of charged side chains of two protein model systems which can be, in part, representative for different molecular sizes and interactivity patterns. One of the two protein is human ubiquitin, hUBQ, considered among the most social ones [8], a behavior which has been ascribed to the high content of charged residues [9]. The other protein is a chitinolytic enzyme from *Ostrinia furnacalis*, oCE, which most likely operates its catalytic role as a dimer, but in the absence of other protein partners [10]. In the crystal structure of the enzyme, its 572 residues are assembled in two domains to form a large globular shape where 151 charged side chains are present with an amino acid composition which is very similar to the average value found in SwissProt database [11], see Fig. S1.

## 2. Distribution and dynamics of charged side chains

Sequence analysis of hUBQ and oCE confirms the already observed frequent recurrence of adjacent residues bearing opposite electric charges [4]. Moreover, hUBQ and oCE charged dipeptides are predominantly located in secondary structure elements which favor close side chains proximity, see Fig. S2. This sequence based preliminary consideration, prompted a systematic structural search for all short distance interactions among charged residues of the two proteins. To account for side chain and backbone flexibility, such investigation has been carried out by means of MD simulations in explicit water.

### 2.1 Charged side chain dynamics in human ubiquitin

In hUBQ sequence, there are 23 charged side chains, 5 D, 6 E, 1H, 7 K and 4 R, mostly surface exposed, as consistently observed in all the solution and crystal structures available from PDB [12]. Along hUBQ sequence the above underlined KE/EK motif is present only twice, *i.e*. K_33_E_34_ and K_63_E_64_. However, within the protein conformational dynamics many other short distance pairwise interactions among the 23 hUBQ charged residues can be established. In order to define the occurrence of the latter interactions, the relative positions of lysyl NZ, glutamyl CD, aspartyl CG arginyl CZ and histidyl CE1 atoms (see PDB amino acid nomenclature) have been monitored along a 500 ns MD simulation in explicit water. These atoms have been chosen for a quantitative description of D, E, H, K and R reciprocal approaches and their interatomic distance, r_IC_, will be discussed hereafter. Close side chain-side chain interactions have been delineated through r_IC_ profiling such the ones shown in Fig. 1. It must be noted that the presence of r_IC_’s shorter than 0.4 nm among the latter atoms is consistent with the formation of hydrogen bonds, HBs, see Fig. S3, which are fundamental turning points for D, E, H, K and R approaching side chains.

**Figure 1:**
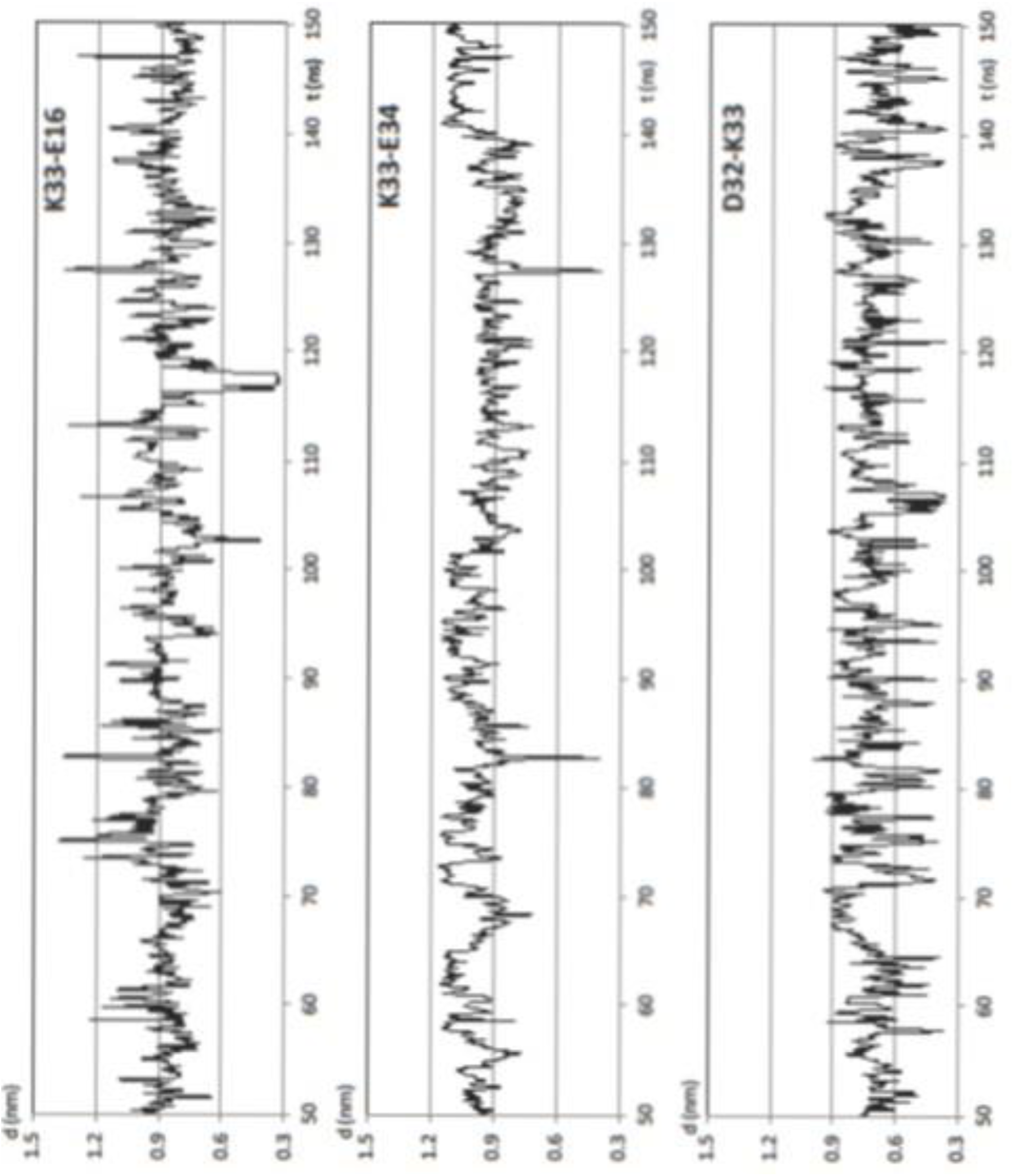
The approach of the NZ atom of hUBQ K33 residue with nearby negatively charged side chains observed in a 100 ns window of the 500 ns MD simulation in explicit water.

Quantitative analysis on charge network dynamics has been performed by sampling r_IC_’s along hUBQ MD trajectory with g_distmap, an *ad hoc* developed GROMACS tool [13], see details given as Supplementary Material. For all charged side chains, their pairwise interactions have been counted to generate r_IC_ binning profiles, such the ones reported in Fig. 2 for K33. Relevant close distance interactions among D, E, H, K and R residues have been considered only in the case that at least 5% of calculated r_IC_’s were below 0.6 nm. Under this limiting condition, from the total 23 × 23 r_IC_ matrix, a reduced set of 32 pairwise interactions was obtained. As shown in Figs. S4, r_IC_ binning profiles have been generated for all short distance approaches. It is interesting to note that all approaches involving side chains bearing opposite charges, consistently with HB formation, exhibit r_IC_ profiles with one sharp peak centered at a distance lower than 0.4 nm.

**Figure 2:**
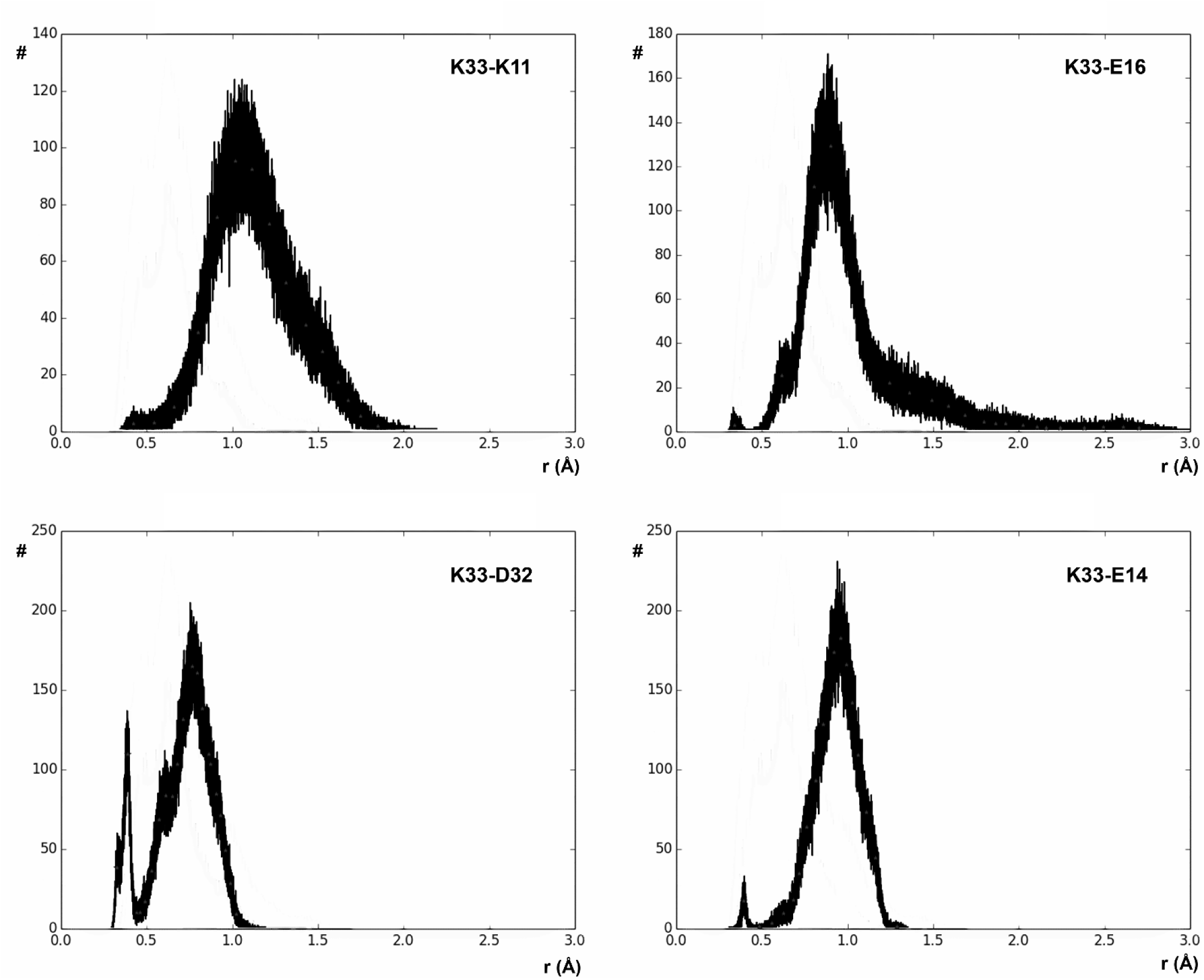
Binned values of r_IC_’s distances probing the approach between charged side chains along hUBQ MD trajectory.

The complete interaction network among hUBQ charged side chains is shown in Fig. 3. It is apparent that all side chains bearing opposite charges establish one or more interactions. Moreover, it can be noted that in the presence of multiple short distance approaches, *e.g*. in the case of the most promiscuous K27 side chain, also equal electric charges are found in close proximity. By observing the evolution of interatomic distances between HB donor and acceptor moieties along the MD trajectory, it is interesting to note that only one HB is present at each time, as in the case of 33 NZ where the amino group jumps from one HB acceptor to another, see Fig. 1.

**Figure 3:**
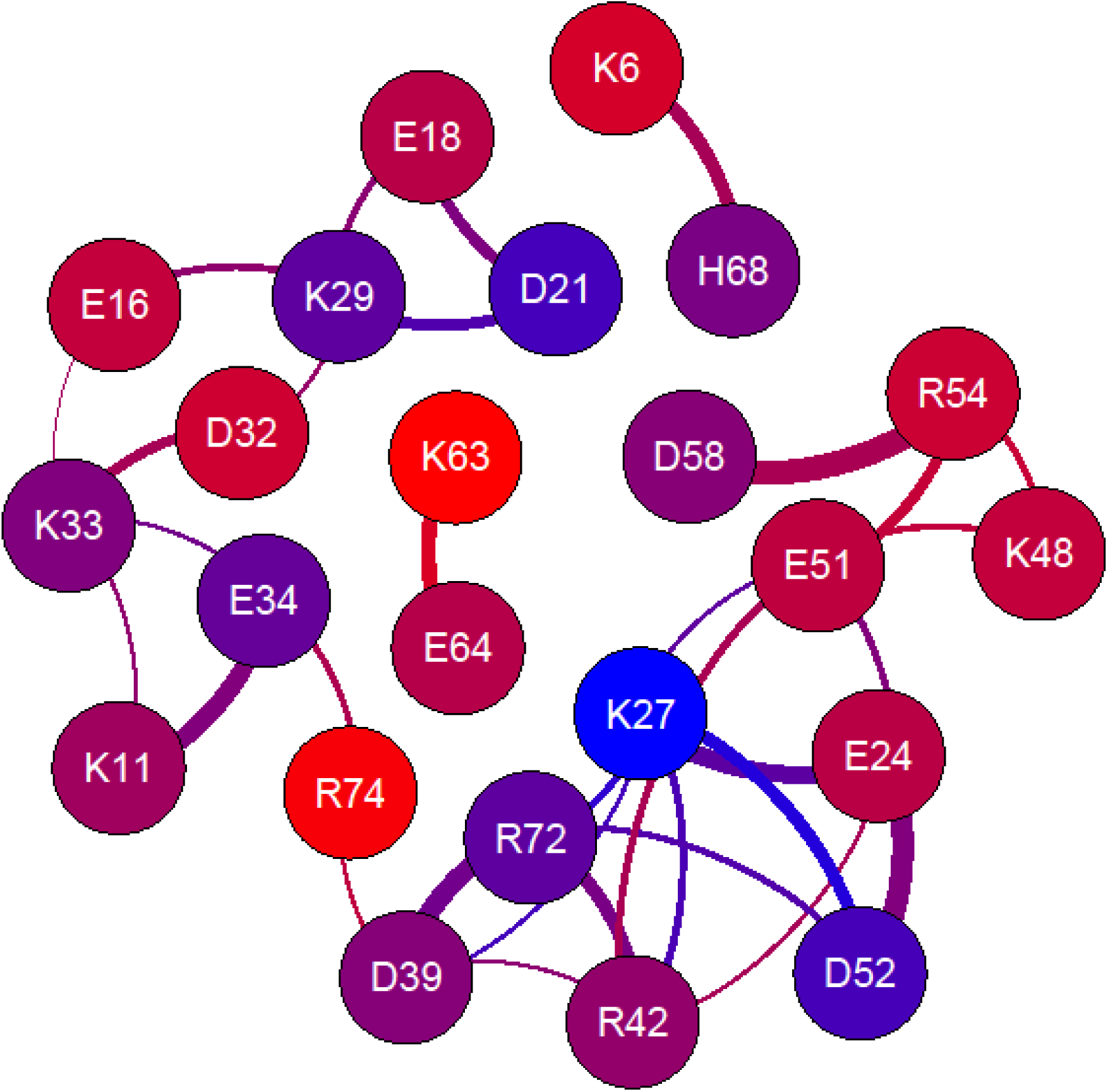
The electric charge interaction network of human ubiquitin. Nodes, representing all charged residues, are colored according to the atom depth values reported in Table SI; edge thickness is proportional to average proximity.

Thus, it can be claimed that the interaction network summarized in Fig. 3 arises from a summation of individual events in rapid succession with a time scale which can be estimated from NMR relaxation studies. Indeed, order parameters, S^2^, of hUBQ lysyl ^15^NZ atoms have been calculated, as well as rotational correlation times accounting for χ_5_ internal side chain rotations [14], showing that the various lysyl side chains experience very different internal dynamics. Experimental data is in very good agreement with our MD results, as the highest S^2^ value determined for the buried K27 NZ atom is consistent with the very limited population of conformers where K27 NZ atom is free from HBs with the neighboring E24, D39, E51 and D52 carboxyl acceptors, see Figs. S4 (panels 16, 24, 25, 26). On the other side, NZ nitrogens of K11, K33, K48 and K63 exhibit i) the lowest S^2^ values, ii) the largest accessible surface areas, see Table SI, and iii) predominantly r_IC_’s values larger than 0.6 nm, see Fig. 3 and Figs. S4. From the same ^15^N NMR relaxation study lysyl χ_5_ reorientations were calculated to range from 25 to 341 ps, respectively for K63 and K27, yielding information on the mobility time scale of the latter hUBQ side chains.

### 2.2 *Charged side chain dynamics in* Ostrinia furnacalis *chitinolytic enzyme*

The same procedure applied to hUBQ has been used to investigate on the electric charge network of oCE. From the crystal structure of the protein, PDB ID code 3NSM, lacking the N terminal 22 amino acids, the relative positions of 41 D, 38 E, 9 H, 35 K and 28 R residues are obtained. In Table SII, depth indexes and DSSP assignments are given for all lysyl NZ, glutamyl CD, aspartyl CG, histidyl CE1 and arginyl CZ atoms of oCE. Most of these 151 residues are distributed on the 592.77 nm^2^ protein surface, almost eight times larger than the one of hUBQ, with time dependent spatial proximities which have been predicted by 30 ns MD simulation in explicit solvent.

From the complex interaction network shown in Fig. 4, derived from the r_IC_ profiles reported in Figs. S5, several features need to be underlined. Differently from hUBQ, eight charged residues are not involved in short distance interactions with other charges. Among these residues, listed with null values of the interaction ranking reported in Table SII, it is interesting to note that R154, K323 and D358 side chains are directly involved in the binding with NAG and TMG-chitotriomycin in the enzyme-inhibitor complex (PDB ID code: 3VTR), suggesting that unperturbed charge density is required by these protein hot spots. The other 143 charged residues are involved in 203 side chain-side chain interactions with ranking values up to 7 for the most social cases offered by the deeply buried H246 and H303 residues, see Table SII. Similarly to what has been found for hUBQ, close distance approaches are mainly due to opposite charges, 137 (67.6 %), but 33 positive to positive and negative to negative encounters (16.2 % each) are found below 0.6 nm. As shown in Fig. 4, oCE 151 charged side chains can be grouped into different patches which can define the boundaries of their concerted motions.

**Figure 4:**
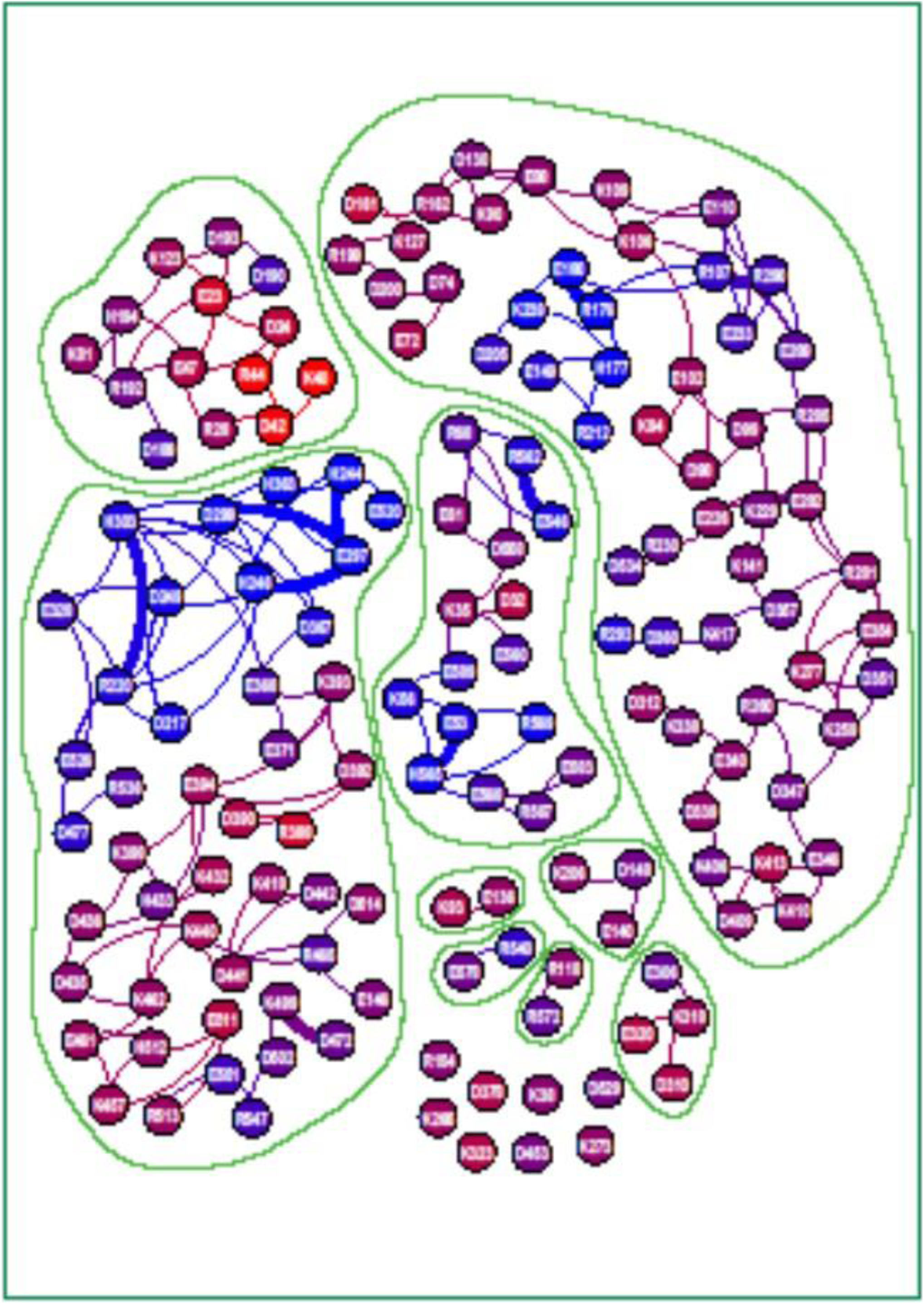
Charged side chain interaction network of *Ostrinia furnacalis* chitinolytic enzyme. Different interaction patches can be identified. Nodes and edges of the graph are shown as already described in Fig. 3 caption. Table SII summarizes some of the parameters underneath the graph.

## 3. From electrostatics to electrodynamics

It is apparent that simple contemplation of the available PDB files for hUBQ and oCE would not yield the wealth of information contained in Figs. 3 and 4. Only MD simulations, indeed, could unveil the complex network of approaches involving D, E, H, K and R side chains of our model proteins, clearly indicating that, along the rapid conformational exchange, electric charges neutralize each other or, conversely, reinforce their strength, as frequently happens in hUBQ and oCE, see one example in Fig. 5. It should be noted also that some of the few oCE charged residues, isolated from other charged ones, bear constant electric messages. This finding is probably required for unambiguous ligand recruitment, as in the case of oCE K323, located at the rim of the active site with its lone positive charge, efficiently orienting the substrate towards binding, see Fig. S6.

**Figure 5:**
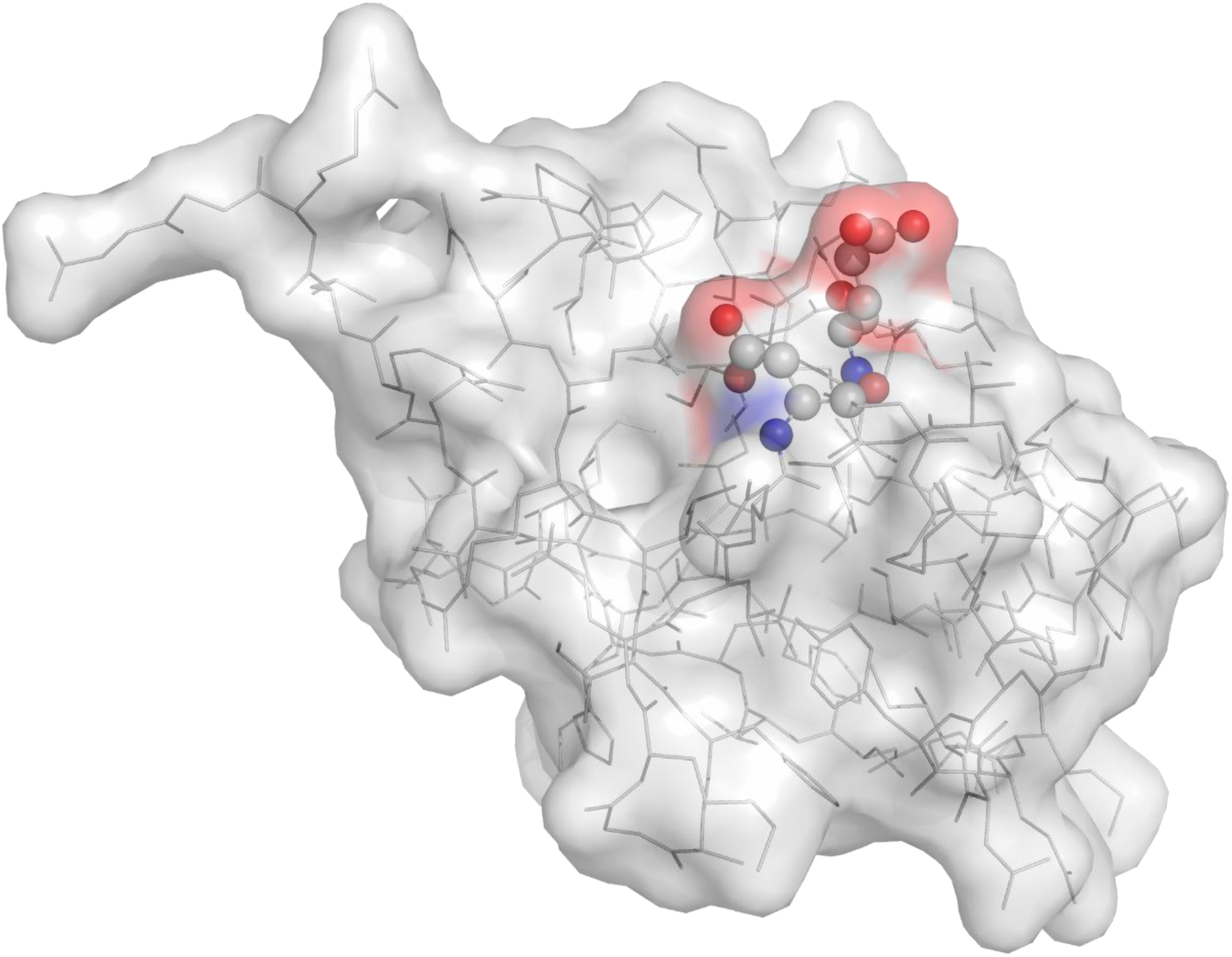
The structural constraint for the short distance E24-D52 interaction in hUBQ.

The above observations delineate a very dynamic panorama for the electric properties of proteins, indicating that electrodynamics rather than electrostatics should be taken into account to predict interactions of proteins with the molecular partners they require for biological activity. This conclusion is of fundamental relevance in the field of protein engineering and drug design, as it can be suggested by the electrostatic analysis calculated for single MD trajectory frames of hUBQ and oCE, as surface flexibility causes, at the same time, changes of molecular shape and electric potential, see Fig. 6. However, an even more important issue must be considered as a consequence of the present analysis on protein charge network. From the macroscopic world, indeed, it is well known that electromagnetic fields are generated by moving charges. In the case that the reciprocal approach of differently charged particles is mainly random, complete averaging out of electromagnetic effects should be expected, but our report indicates that this is not the case for the electric charge assembly presented by folded proteins. Indeed, side chain flexibility, in the presence of multiple attractive or repulsive electric interactions, yields highly concerted motions, where reduction or even absence of averaging effects has to be expected.

**Figure 6:**
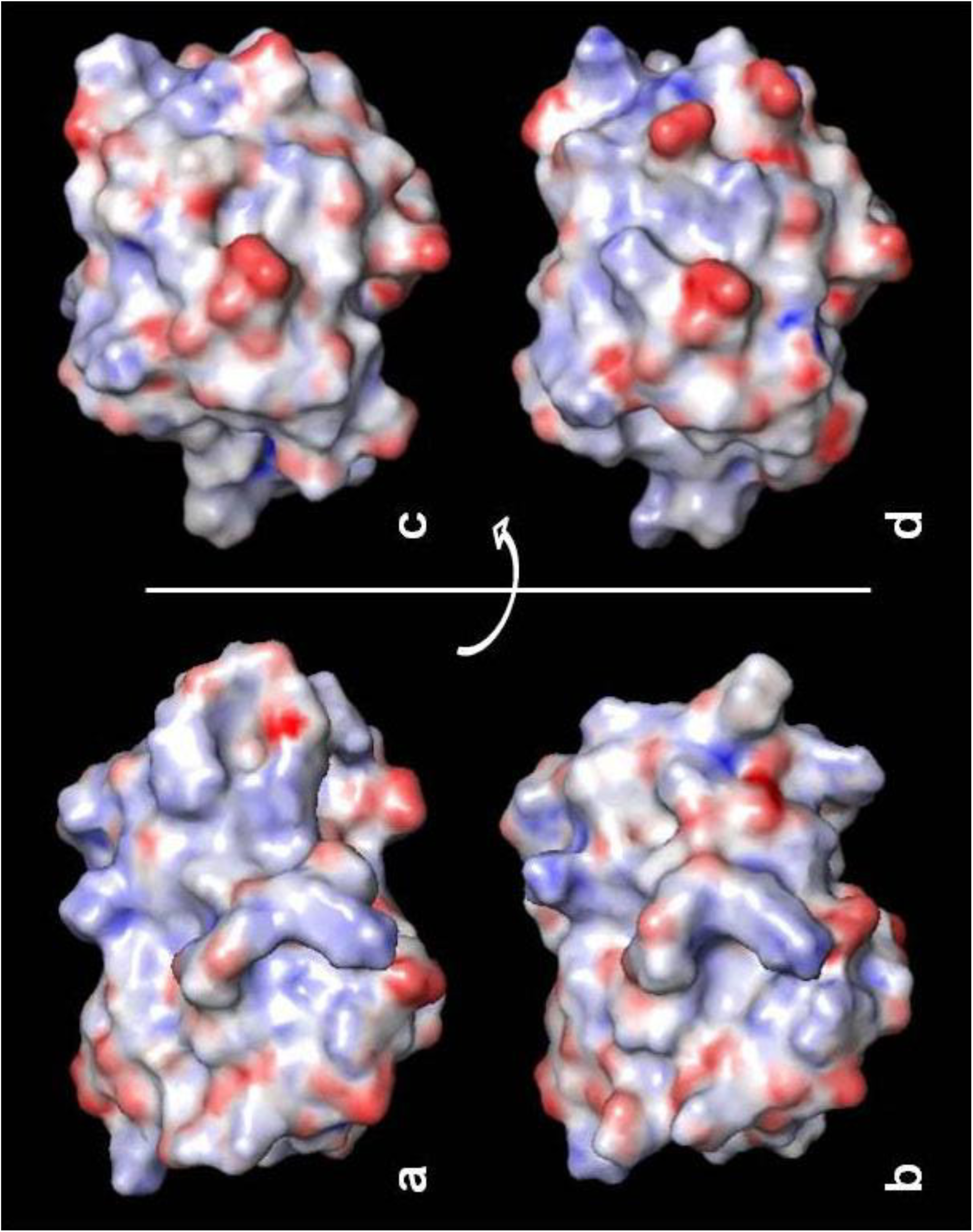
Protein Connolly surfaces from two different snapshots of MD simulation of human ubiquitin, panel a/b and c/d. Surfaces are colored according to the projection of electrostatic maps obtained with Pymol APBS plugin. Animated view of hUBQ MD simulation is available as Supplementary Material.

NMR studies indicate that changes of charged orientations for lysyl side chains occur in a picosecond time scale, suggesting a time scale for the electric interactions among protein charges. It follows that electromagnetic fields in the terahertz frequency region should be generated by each element of the charge interaction network. In this respect, it must be pointed out that different surface exposures and HB involvements of approaching charged partners determine characteristic reorientational freedom and, hence, electromagnetic fields.

Thus, local orientations and distance fluctuations of each charge-charge interaction are deeply encoded in protein structural dynamics yielding a highly specific electromagnetic fingerprinting with D, E, H, K and R side chains acting as protein antennas to transmit and receive long distance messages with their molecular partners. The electrodynamics associated to protein charge interaction network should be analyzed in details for defining all the features of this radiative transfer of molecular information. Terahertz spectroscopy [15,16], with systematic investigations on the functional effects of charged amino acid substitutions to specific interacting protein systems, e.g. with E/Q and D/N replacements, is a promising experimental approach for supporting our hypothesis of long distance protein-protein communications. Surface Plasmon Resonance [17] could be also a powerful technique to delineate the effects of modified charge networks in protein-protein interactions by analyzing the corresponding k_on_ rate constants.

In any case, even though direct evidence of these electromagnetic effects might be difficult to be experimentally detected, all the ingredients are there, as the presence of charged amino acid side chains coherently reorienting on protein surfaces just supports the present hypothesis: a Coulomb’s egg.

## 4. Methods

500 ns and 30 ns MD simulations respectively for hUQB and oCE have been performed by using GROMACS package [13] on 1UBQ and 3NSM PDB structures, solvated in a cubic box of equilibrated TIP3P water molecules [18]. The time dependent evolution along the MD trajectories of all the approaches among charged side chains has been monitored by calculating in each of the MD frames, sampling interval of 1 ps, the distance, r_IC_, between lysyl NZ, glutamyl CD, aspartyl CG arginyl CZ and histidyl CE1 atoms. These atoms have been selected as representative ones to assign electric charge positions irrespective of actual electric charge distribution. Upon binning of r_IC_ distances determined along the MD trajectory, all the possible combinations among the latter atoms were analyzed with a new GROMACS tool, g_distmap, which has two groups of atoms as input. Let group 1 include atoms A, B, C, D and group 2 include atoms E, F, G, g_distmap calculates, along the MD trajectory, all the distances AE, AF, AG, BE, BF etc. For each atom pair a n-dimensional XVG file can be obtained where distances and simulation times are coupled. Then, only those side chain interactions having a number of distances higher than 5 % of r_IC_’s < 0.6 nm have been selected for further analysis. Charged side chain networks have been analyzed and visualized by using Gephi 0.8.2-beta software [19]. PyMOL v 1.7 has been used for all molecular graphics and, through the APBS plugin [20], for electrostatics calculations on each MD frames. Atom depth indexes, D_i_, have been calculated by using SADIC algorithm, as explained in the Supplementary Material.

## Acknowledgements

This work has received financial support from the Istituto Toscano Tumori. Thanks are due to Profs. Piero Andrea Temussi and Michael McCormick for precious suggestions.

## Appendix A. Supplementary data

Supplementary data associated with this article can be found, in the online version, at http://dx.doi.org/xxxxx

