## Supplemental Tables and Figures for "Long range electromagnetic effects drive protein-protein approaches: an egg of Coulomb"

#### Atom depth calculations

Atom depth is considered among the structural descriptors that correlate protein structure with folding and functional properties. The distance between an atom and the nearest water molecule or the closest surface dot has been proposed as a measure of the atom depth, but, in both cases, the three-dimensional character of depth is lost. To account for the overall molecular shape in atom depth calculations, measurements of intersections between the molecular volume and a sphere of a suitable radius, centered on the atom whose depth has to be quantified, have been proposed by the authors in a previous paper [1]. It is apparent that the smaller is the exposed volume, the deeper is the three-dimensional insertion of the investigated atom inside the molecular structure.

As, in general, depth is a very relative term which depends on the overall size and shape of the object under discussion, atom depth indexes, $D_i$'s are suggested as useful parameters to compare atom insertions within the considered molecular systems. Thus, for an atom i of a given molecule and a sampling radius r, a depth index $D_{i,r}$ may be defined as

$$D_{i,r} = 2V_{i,r}/V_{0,r}$$

where $V_{i,r}$ is the exposed volume of a sphere of radius $r$ centered on atom $i$ and $V_{0,r}$ is the exposed volume of the same sphere when centered on an isolated atom. Thus, the limiting $D_i$ values 0 and 2 are obtained for a totally buried or totally isolated atom, respectively.

---

[1] D. Varrazzo, A. Bernini, O. Spiga, A. Ciutti, S. Chiellini, V. Venditti, L. Bracci, N. Niccolai, Three-dimensional computation of atom depth in complex molecular structures, Bioinformatics 21 (2005) 2856-2860.

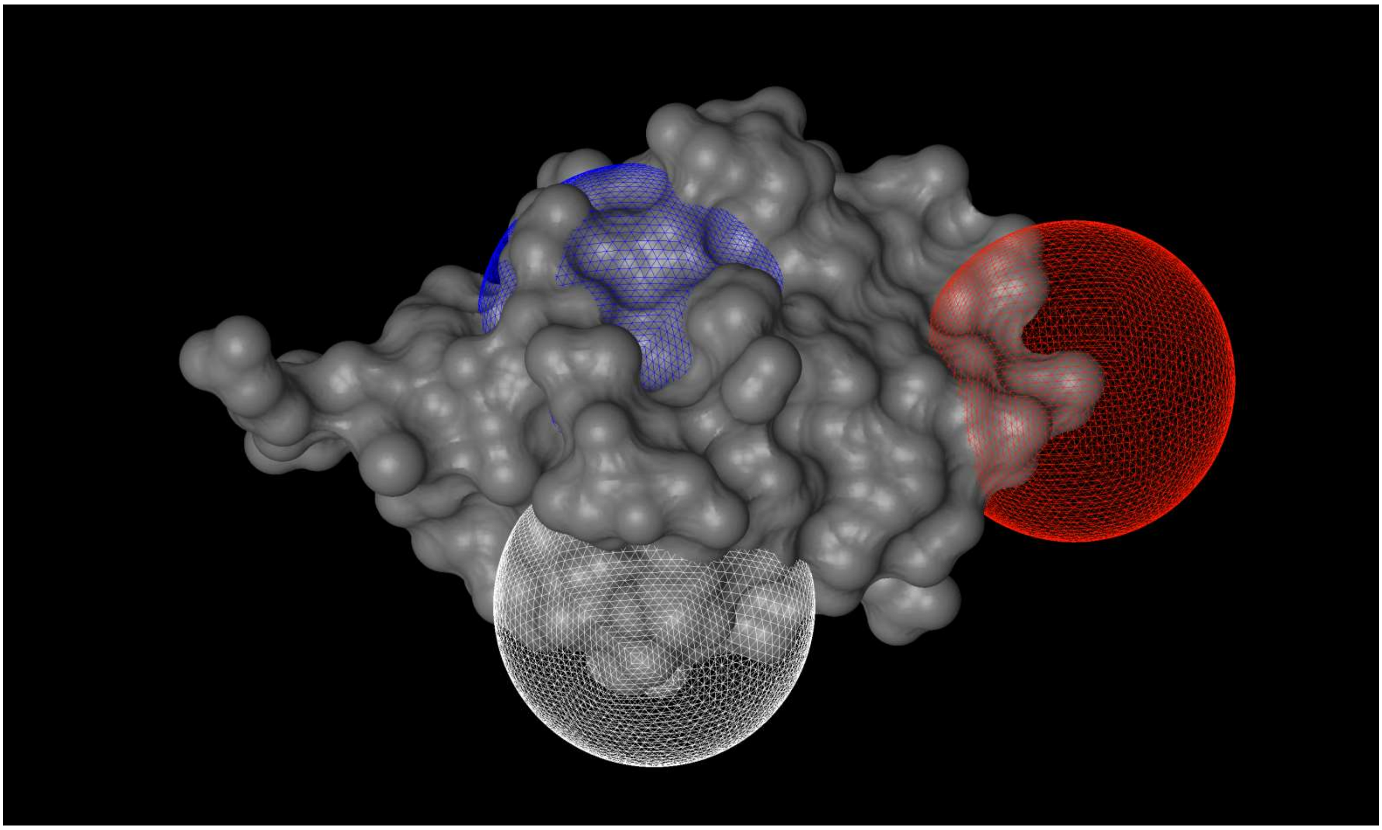

The exposed volumes of L43 CA (buried, color blue, $D_i$ = 0.05), E34 CD (intermediate, color white, $D_i$ = 0.76), K63 NZ (exposed, color red, $D_i$ = 1.44) are shown as mesh on Ubiquitin surface (PDB: 1UBQ) by drawing a sphere of 9 Å radius on each atom.

The algorithm for the calculation of $D_i$ , extensively described in [1], has been implemented in the software Simple Atom Depth Index Calculator (SADIC), available at http://www.sbl.unisi.it/prococoa. The software outputs the $D_i$ values calculated for the protein structure given in PDB format in the B-factor field of the same PDB file, allowing for coloring by depth index with the most common graphical viewers, like PyMol (http://www.pymol.org).

### Table SI

| Atom | # interactions | $D_i$ | DSSP |
|---|---|---|---|
| NZ K6 | 1 | 1.29 | E |
| NZ K11 | 2 | 1.06 | |
| CD E16 | 2 | 1.22 | E |
| CD E18 | 2 | 1.16 | |
| CG D21 | 2 | 0.60 | S |
| CD E24 | 3 | 1.17 | H |
| NZ K27 | 6 | 0.35 | H |
| NZ K29 | 4 | 0.77 | H |
| CG D32 | 2 | 1.23 | H |
| NZ K33 | 4 | 0.93 | H |
| CD E34 | 3 | 0.76 | H |
| CG D39 | 4 | 0.90 | G |
| CZ R42 | 5 | 1.00 | E |
| NZ K48 | 2 | 1.18 | E |
| CD E51 | 5 | 1.19 | |
| CG D52 | 4 | 0.66 | T |
| CZ R54 | 3 | 1.21 | S |
| CG D58 | 1 | 0.92 | G |
| NZ K63 | 1 | 1.44 | T |
| CD E64 | 1 | 1.10 | T |
| CE1 H68 | 1 | 0.91 | E |
| CZ R72 | 4 | 0.75 | |
| CZ R74 | 2 | 1.42 | S |

### Table SII

| Atom | # interactions | $D_i$ | DSSP |
|---|---|---|---|
| CD E23 | 6 | 1.55 | |
| CG D24 | 3 | 1.33 | |
| CZ R28 | 2 | 0.96 | E |
| CG D32 | 1 | 1.30 | E |
| NZ K35 | 4 | 1.17 | E |
| NZ K38 | 0 | 1.00 | E |
| NZ K40 | 1 | 1.42 | E |
| CG D42 | 3 | 1.30 | |
| CZ R44 | 2 | 1.77 | S |
| CD E47 | 5 | 1.17 | S |
| CD E53 | 3 | 0.19 | H |
| NZ K56 | 3 | 0.21 | H |
| CD E61 | 2 | 0.93 | T |
| CZ R68 | 4 | 0.66 | |
| CD E72 | 1 | 1.24 | |
| CG D74 | 2 | 1.03 | E |
| NZ K81 | 2 | 1.10 | E |
| CD E88 | 5 | 1.10 | E |
| NZ K90 | 3 | 1.16 | E |
| NZ K93 | 1 | 1.25 | S |
| NZ K94 | 2 | 1.43 | |
| CG D98 | 3 | 1.10 | H |
| CG D99 | 4 | 1.00 | H |
| CD E102 | 4 | 1.10 | H |
| NZ K106 | 5 | 1.28 | H |
| CZ R107 | 6 | 0.43 | T |
| NZ K109 | 3 | 1.00 | H |
| CD E110 | 5 | 0.90 | H |
| CZ R118 | 1 | 1.10 | T |
| NZ K123 | 3 | 1.17 | |
| NZ K127 | 2 | 0.99 | E |
| CG D130 | 3 | 0.87 | E |
| CD E136 | 1 | 1.11 | S |
| CD E140 | 1 | 1.01 | |
| NZ K141 | 2 | 0.98 | |
| CD E146 | 1 | 1.03 | T |
| CG D148 | 2 | 0.83 | |
| CD E149 | 2 | 0.22 | |
| CZ R154 | 0 | 1.06 | E |
| CG D161 | 2 | 1.38 | T |
| CZ R162 | 4 | 1.11 | E |
| CZ R176 | 3 | 0.05 | H |
| CE1 H177 | 4 | 0.05 | H |
| CD E180 | 3 | 0.07 | H |
| CG D189 | 1 | 0.54 | E |
| CG D190 | 2 | 0.51 | T |
| CZ R192 | 5 | 0.78 | T |
| CG D193 | 3 | 0.96 | T |
| CE1 H194 | 4 | 0.95 | E |
| CZ R199 | 2 | 1.13 | S |
| CG D200 | 2 | 0.96 | S |
| CG D205 | 1 | 0.37 | E |
| NZ K206 | 1 | 1.19 | |
| CZ R212 | 2 | 0.13 | E |
| CG D217 | 3 | 0.14 | E |
| CZ R220 | 6 | 0.23 | S |
| CD E226 | 3 | 1.15 | H |
| NZ K229 | 2 | 0.94 | H |
| CZ R230 | 2 | 0.74 | H |
| CD E233 | 3 | 0.51 | H |
| NZ K239 | 3 | 0.03 | T |
| CE1 H244 | 4 | 0.02 | E |
| CE1 H246 | 7 | 0.06 | E |
| CG D249 | 6 | 0.14 | |
| NZ K259 | 5 | 1.05 | S |
| CZ R260 | 3 | 0.89 | S |
| NZ K265 | 0 | 1.20 | H |
| NZ K273 | 0 | 1.15 | S |
| NZ K277 | 4 | 1.18 | H |
| CZ R281 | 5 | 1.10 | H |
| CD E282 | 4 | 1.08 | H |
| CZ R285 | 4 | 0.87 | H |
| CD E289 | 4 | 0.68 | H |
| CZ R290 | 4 | 0.49 | T |
| CZ R293 | 1 | 0.22 | E |
| CD E297 | 5 | 0.04 | E |
| CG D299 | 6 | 0.06 | E |
| CE1 H303 | 8 | 0.22 | S |
| CD E306 | 1 | 0.77 | T |
| CG D310 | 1 | 1.30 | T |
| CG D312 | 1 | 1.11 | |
| NZ K318 | 3 | 1.24 | T |
| CD E320 | 1 | 1.30 | S |
| NZ K323 | 0 | 1.34 | G |
| CD E328 | 4 | 0.43 | S |
| NZ K338 | 2 | 1.01 | |
| CG D339 | 2 | 1.09 | T |
| CD E340 | 3 | 1.05 | H |
| CD E346 | 3 | 0.95 | H |
| CG D347 | 3 | 0.80 | H |
| CG D351 | 3 | 0.65 | H |
| CD E354 | 4 | 1.04 | H |
| CG D357 | 3 | 0.77 | |
| CG D360 | 2 | 0.45 | S |
| CE1 H363 | 3 | 0.04 | E |
| CG D367 | 3 | 0.18 | |
| CD E368 | 4 | 0.56 | |
| CD E371 | 2 | 0.83 | H |
| CG D378 | 0 | 1.25 | H |
| CZ R388 | 1 | 1.57 | T |
| CG D390 | 2 | 1.29 | |
| CG D392 | 3 | 1.21 | S |
| NZ K393 | 4 | 1.19 | S |
| CD E394 | 5 | 1.18 | S |
| NZ K398 | 3 | 1.05 | H |
| NZ K406 | 3 | 0.93 | H |
| CG D409 | 3 | 0.97 | H |
| NZ K410 | 4 | 0.98 | H |
| NZ K413 | 3 | 1.36 | H |
| NZ K417 | 2 | 0.94 | S |
| NZ K418 | 2 | 1.12 | |
| NZ K432 | 3 | 1.27 | T |
| CE1 H433 | 4 | 0.81 | T |
| CG D435 | 3 | 0.97 | H |
| CG D436 | 3 | 1.08 | H |
| NZ K440 | 4 | 1.21 | T |
| CG D441 | 4 | 1.15 | T |
| CG D442 | 2 | 0.88 | T |
| CG D453 | 0 | 0.91 | |
| NZ K457 | 3 | 1.29 | H |
| CD E461 | 3 | 1.16 | H |
| NZ K462 | 4 | 1.16 | T |
| CZ R465 | 4 | 0.64 | E |
| CG D472 | 1 | 0.74 | H |
| CG D477 | 2 | 0.26 | T |
| NZ K499 | 2 | 0.91 | H |
| CG D502 | 2 | 0.80 | H |
| CD E511 | 3 | 1.31 | G |
| CE1 H512 | 3 | 1.10 | G |
| CZ R513 | 2 | 1.22 | G |
| CG D514 | 1 | 1.02 | G |
| CD E520 | 1 | 0.01 | E |
| CD E526 | 3 | 0.36 | T |
| CG D529 | 0 | 0.88 | |
| CG D534 | 1 | 0.57 | H |
| CZ R536 | 1 | 0.64 | H |
| CZ R540 | 1 | 0.34 | H |
| CD E546 | 2 | 0.18 | H |
| CZ R547 | 2 | 0.52 | H |
| CD E551 | 2 | 0.54 | |
| CG D558 | 3 | 0.95 | G |
| CD E560 | 1 | 0.58 | H |
| CZ R562 | 2 | 0.25 | H |
| CE1 H565 | 4 | 0.10 | H |
| CZ R567 | 2 | 0.75 | H |
| CD E568 | 3 | 0.53 | H |
| CZ R569 | 2 | 0.13 | H |
| CZ R572 | 1 | 0.77 | H |
| CD E578 | 1 | 0.82 | |
| CD E583 | 2 | 0.84 | H |
| CD E589 | 2 | 0.33 | T |

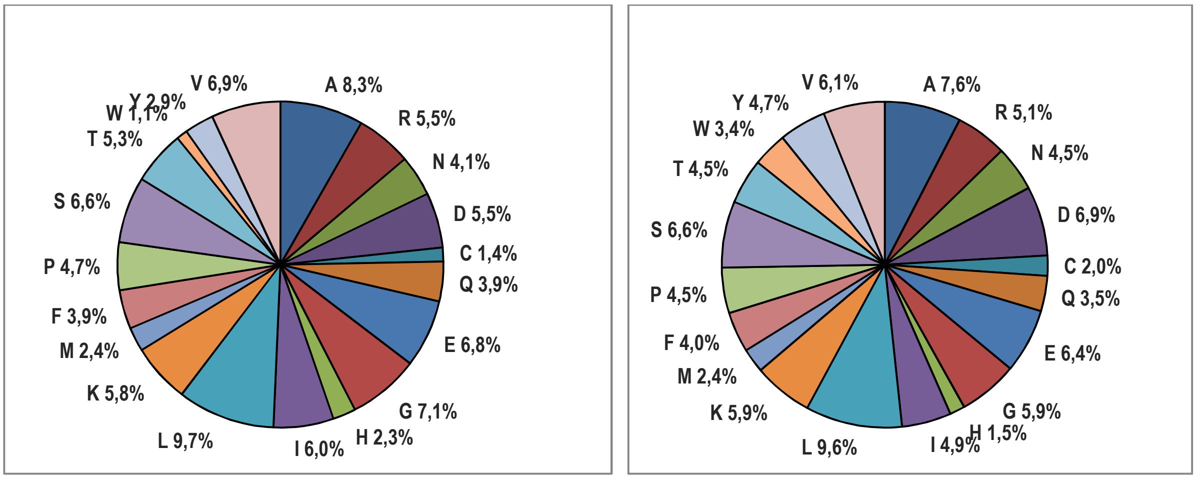


**Figure S1**: left panel: average amino acid composition of proteins contained in release 2014_06 of 11-Jun-14 of UniProtKB/Swiss-Prot (http://web.expasy.org); right panel: amino acid composition of *Ostrinia furnacalis* chitinolytic enzyme (UniProtKB/Swiss-Prot ID: Q06GJ0)

**Human ubiquitin**

MQIFVKTLTGKTITLEVEPSDTIENVKAKIQD$^{33}$**K$_{H}$E$_{H}$**GIPPDQQRLIFAGKQLEDGRTLSDYNIQ$^{63}$**K$_{T}$E$_{T}$**STLHLVLRLRGG

**Ostrinia furnacalis chitinolytic enzyme**

EDVVWRWSCDNGKCVKLKNDPRSSEPALSLEACKMFCNEYGLLWPRPTGEADLGNFLSKINLNSIEVKILKKGATDDLMEA
AAKRF$^{109}$**K$_{H}$E$_{H}$**QVSLAIPRGSTPKLTGKAVDVYLVNENPN$^{140}$**EK**AFSLEMDESYGLRVSPSGA$^{161}$**D$_{T}$R$_{E}$**VNATITANSFFGMRH
GLETLSQLFVFDDI$^{192}$**R$_{T}$D$_{T}$**HLLMV$^{199}$**R$_{S}$D$_{S}$**VNIS$^{205}$**D$_{H}$K$_{H}$**PVYPYRGILLDTARNYYSIESIKRTIEAMAAVKLNTFHWHITDSQS
FPFVTTKRPNLYKFGALSPQKVYTKAAI$^{281}$**R$_{H}$E$_{H}$**VVRFGL$^{289}$**E$_{H}$R$_{H}$**GVRVLPEFDAPAHVGEGWQDTDLTVCFKAEPWKSYCV
EPPCGQLNPT$^{338}$**KD$_{T}$**ELYQYLEDIYSDMAEVFDTTDIFHMGGDEVSEACWNSSDSIQNFMMQNRWDLD$^{393}$**K$_{H}$E$_{H}$**SFLKLW
NYFQQKAQ$^{409}$**D$_{H}$K$_{H}$**AYKAFGKKLPLILWTSTLTNYKHIDDYLN$^{440}$**K$_{T}$D$_{T}$**DYIIQVWTTGVDPQIKGLL$^{461}$**E$_{H}$K$_{T}$**GYRLIMSNYDA
LYFDCGYGAWVGAGNNWCSPYIGWQKVYDNSPAVIALEH$^{513}$**R$_{G}$D$_{G}$**QVLGGEAALWSEQSDTSTLDGRLWPRAAALA$^{546}$**E$_{H}$
R$_{H}$**LWAEPATSWQDAEYRMLHIR$^{568}$**E$_{H}$R$_{H}$**LVRMGIQAESLQPEWCYQNEGYCYS

**Figure S2**: Dipeptide fragments of charged amino acids are highlighted in red; numbers refer to residue position and subscripts define DSSP assignments.

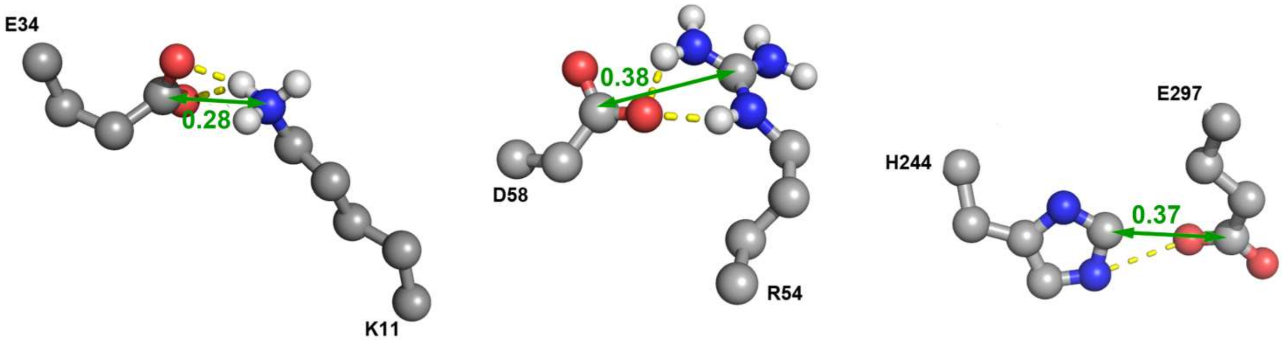


**Figure S3**: Closest approaches of charged side chains stabilized by the formation of hydrogen bonding. Green arrows highlight the distances $r_{IC}$ of E34 CG, D58 CD and E297 CG atoms respectively with K11 NZ, R54 CZ and H244 CE1 in *Ostrinia furnacalis* chitinolytic enzyme.

**Figure S4**: Binned distributions of close distance interactions between D, E, H, K and R side chains in hUBQ as calculated along the 500 ns MD trajectory with g_distmap tool. Only side chain-side chain approaches exhibiting more than 5 % occurrences at $r_{IC} < 0.6$ nm have been considered.

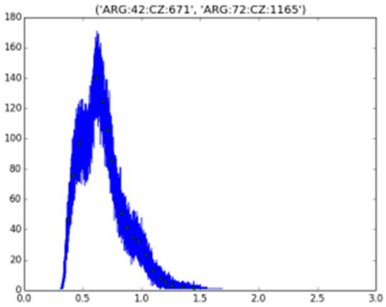


1. R42 CZ – R72 CZ

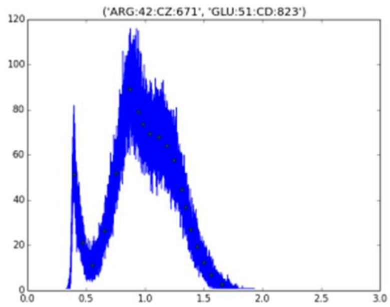


2. R42 CZ – E51 CD

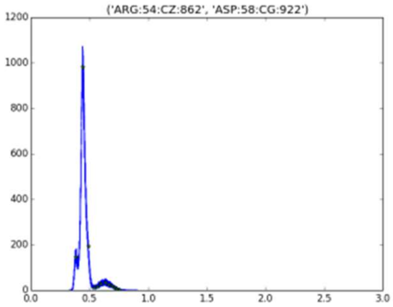


3. R54 CZ – D58 CG

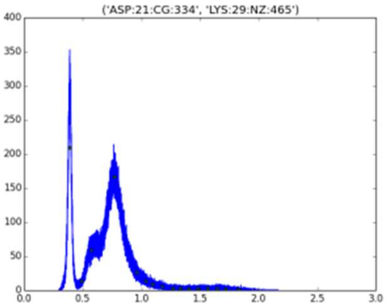


4. D21 CG – K29 NZ

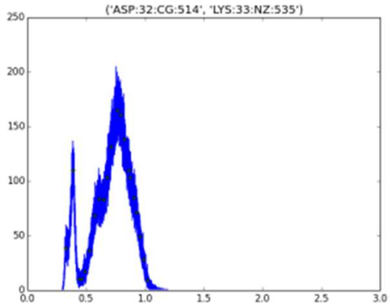


5. D32 CG – K33 NZ

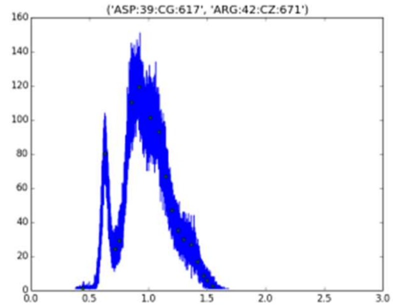


6. D39 CG – R42 CZ

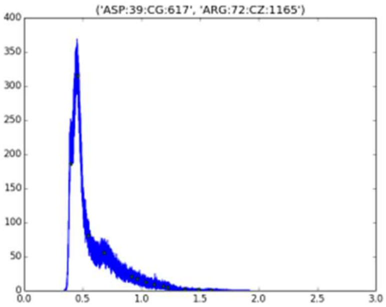


7. D39 CG – R72 CZ

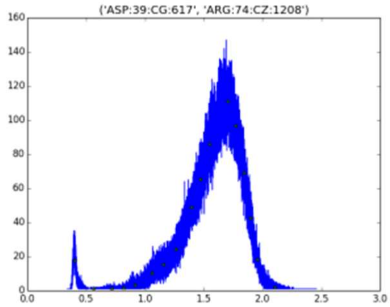


8. D39 CG – R74 CZ

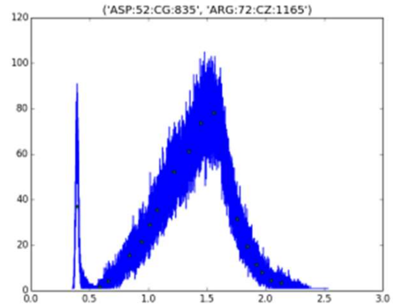


9. D52 CG – R72 CZ

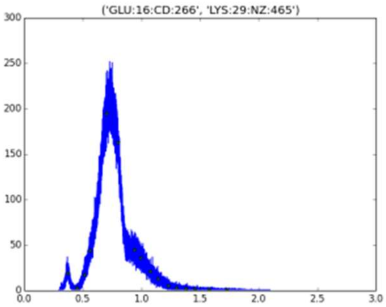


10. E16 CD – K29 NZ

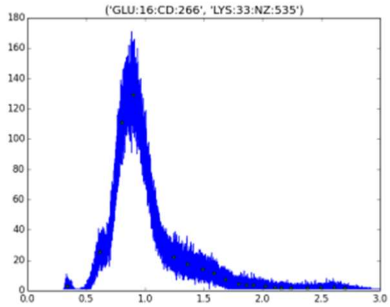


11. E16 CD – K33 NZ

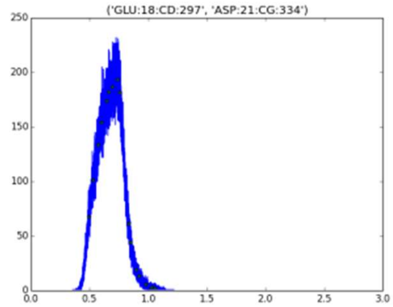


12. E18 CD – D21 CG

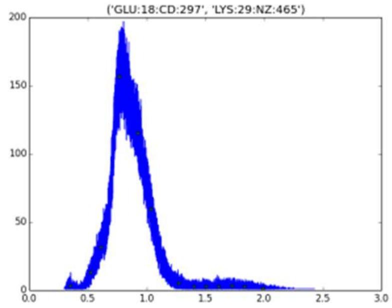


13. E18 CD – K29 NZ

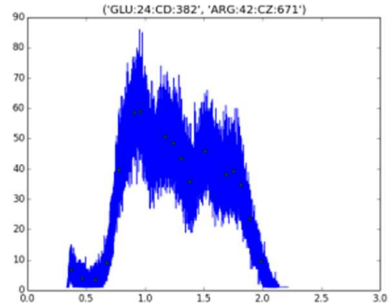


14. E24 CD – R42 CZ

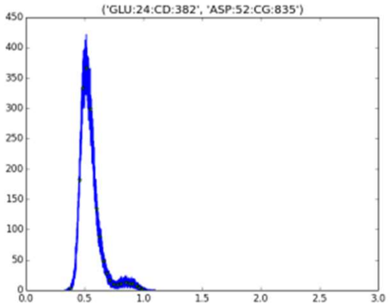


15. E24 CD – D52 CG

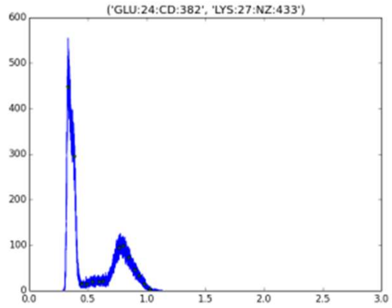


16. E24 CD – K27 NZ

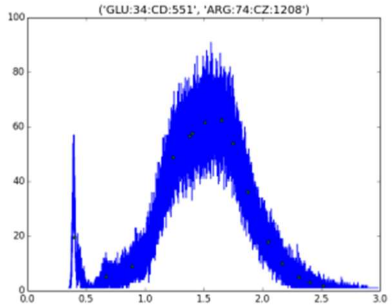


17. E34 CD – R74 CZ

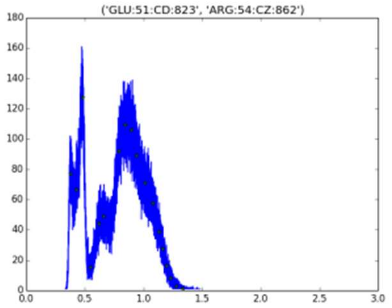


18. E51 CD – R54 CZ

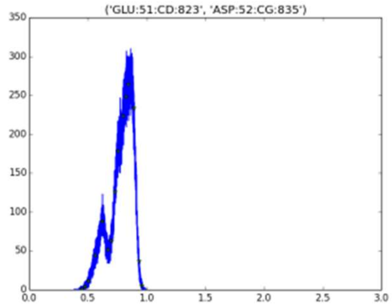


19. E51 CD – D52 CG

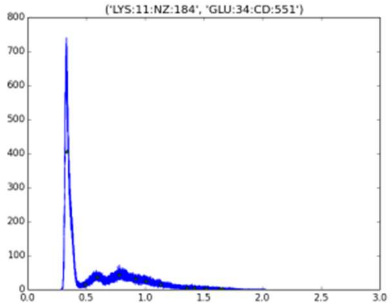


20. K11 NZ – E34 CD

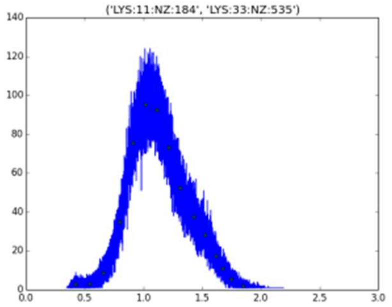


21. K11 NZ – K33 NZ

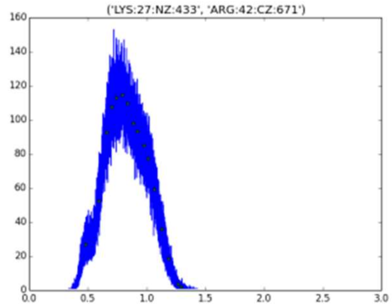


22. K27 NZ – R42 CZ

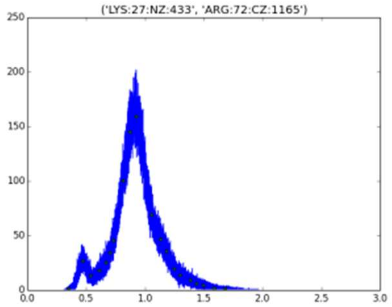


23. K27 NZ – R72 CZ

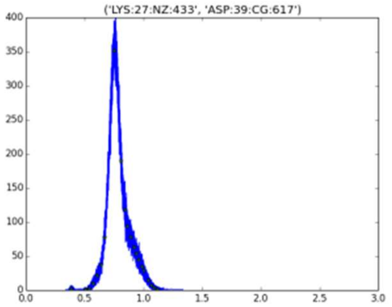


24. K27 NZ – D39 CG

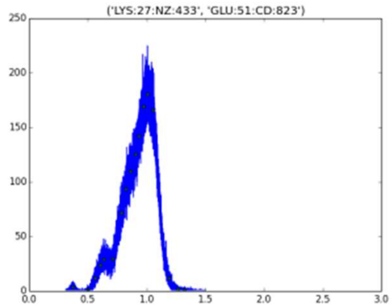


25. K27 NZ – E51 CD

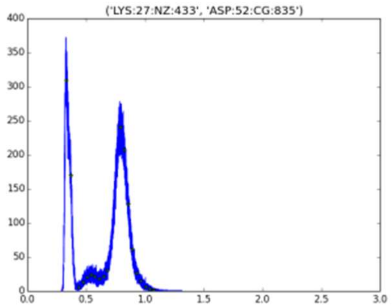


26. K27 NZ – D52 CG

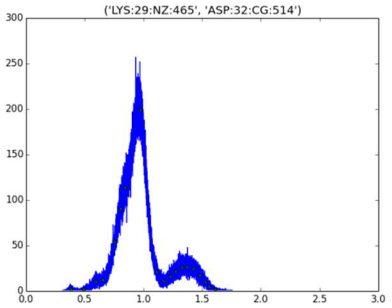


27. K29 NZ – D32 CG

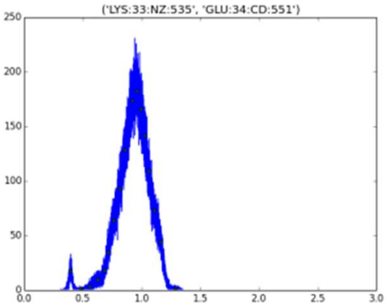


28. K33 NZ – E34 CD

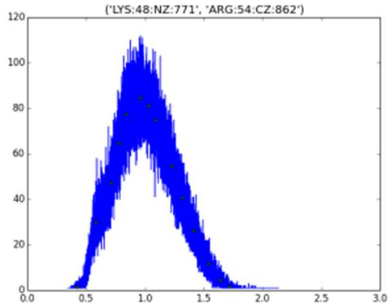


29. K48 NZ - R54 CZ

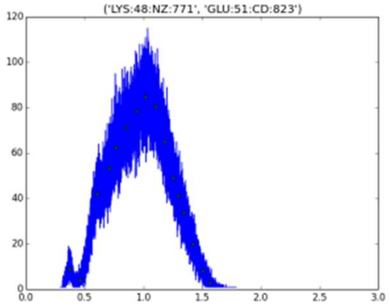


30. K48 NZ – E51 CD

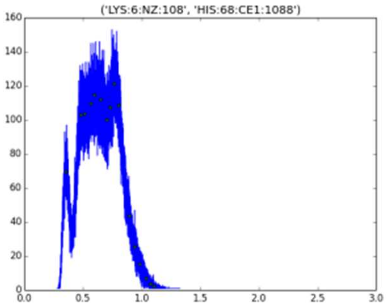


31. K6 NZ – H8 CE1

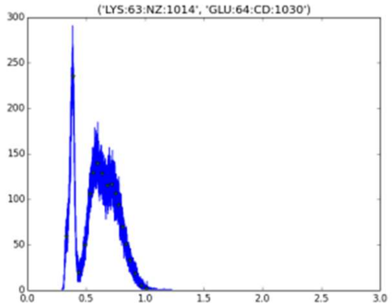


32. K63 NZ – E64 CD

Figure S5: Binned distributions of close distance interactions between D, E, H, K and R side chains in oCE as calculated along the 30 ns MD trajectory with g_distmap tool. Only side chain-side chain approaches exhibiting more than 5 % occurrences at $r_{IC} < 0.6$ nm have been considered.

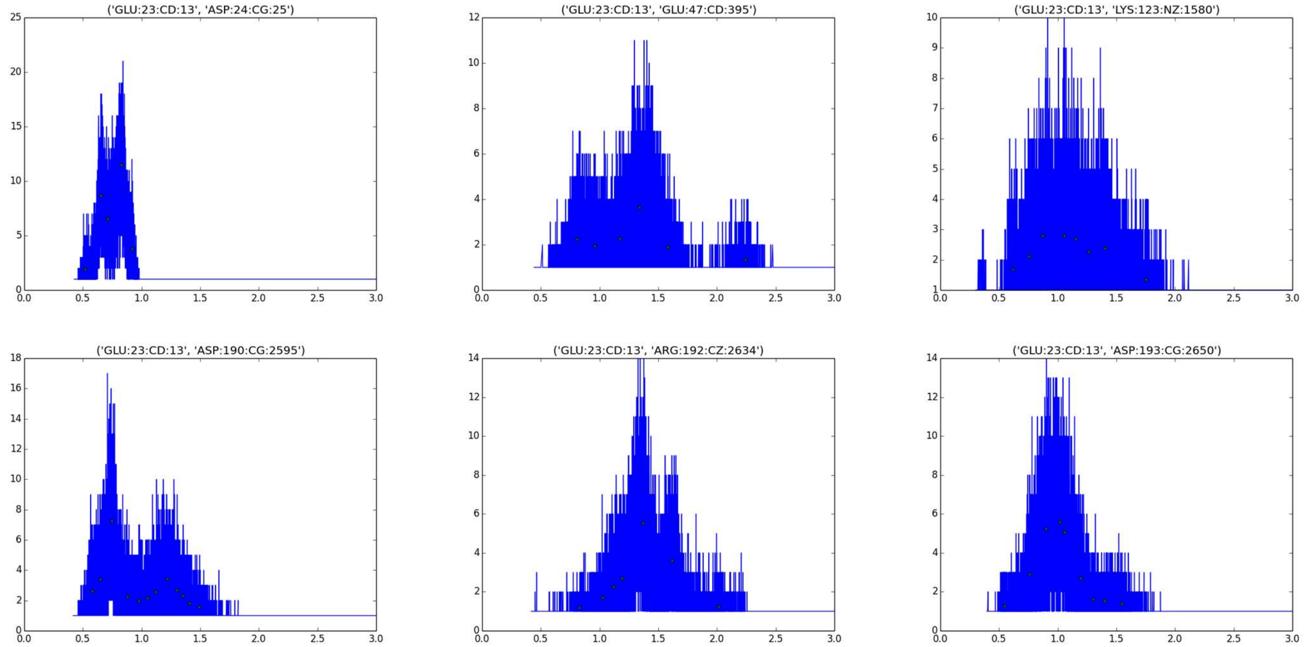

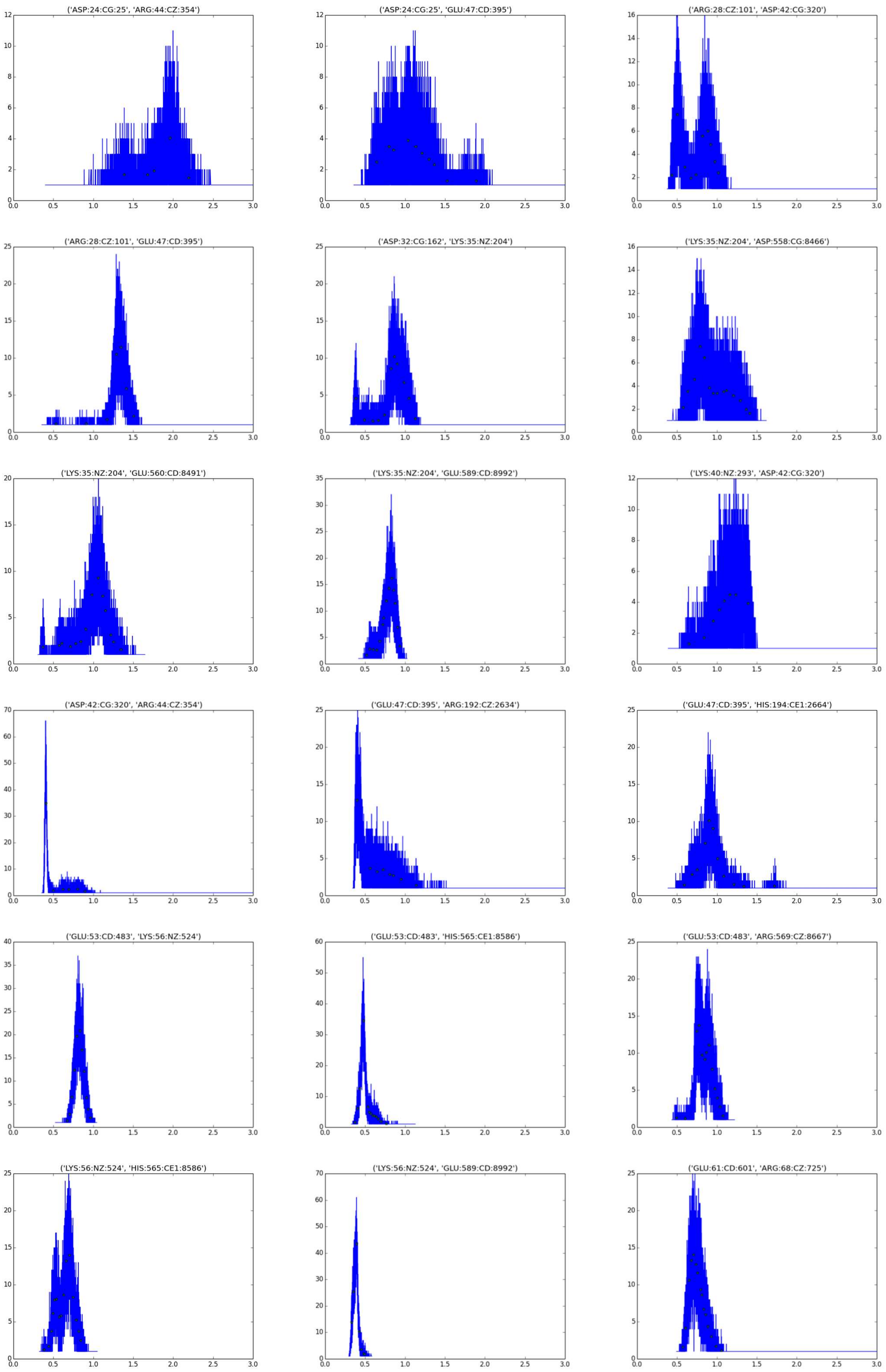
('ASP:24:CG:25', 'ARG:44:CZ:354')
('ASP:24:CG:25', 'GLU:47:CD:395')
('ARG:28:CZ:101', 'ASP:42:CG:320')
('ARG:28:CZ:101', 'GLU:47:CD:395')
('ASP:32:CG:162', 'LYS:35:NZ:204')
('LYS:35:NZ:204', 'ASP:558:CG:8466')
('LYS:35:NZ:204', 'GLU:560:CD:8491')
('LYS:35:NZ:204', 'GLU:589:CD:8992')
('LYS:40:NZ:293', 'ASP:42:CG:320')
('ASP:42:CG:320', 'ARG:44:CZ:354')
('GLU:47:CD:395', 'ARG:192:CZ:2634')
('GLU:47:CD:395', 'HIS:194:CE1:2664')
('GLU:53:CD:483', 'LYS:56:NZ:524')
('GLU:53:CD:483', 'HIS:565:CE1:8586')
('GLU:53:CD:483', 'ARG:569:CZ:8667')
('LYS:56:NZ:524', 'HIS:565:CE1:8586')
('LYS:56:NZ:524', 'GLU:589:CD:8992')
('GLU:61:CD:601', 'ARG:68:CZ:725')

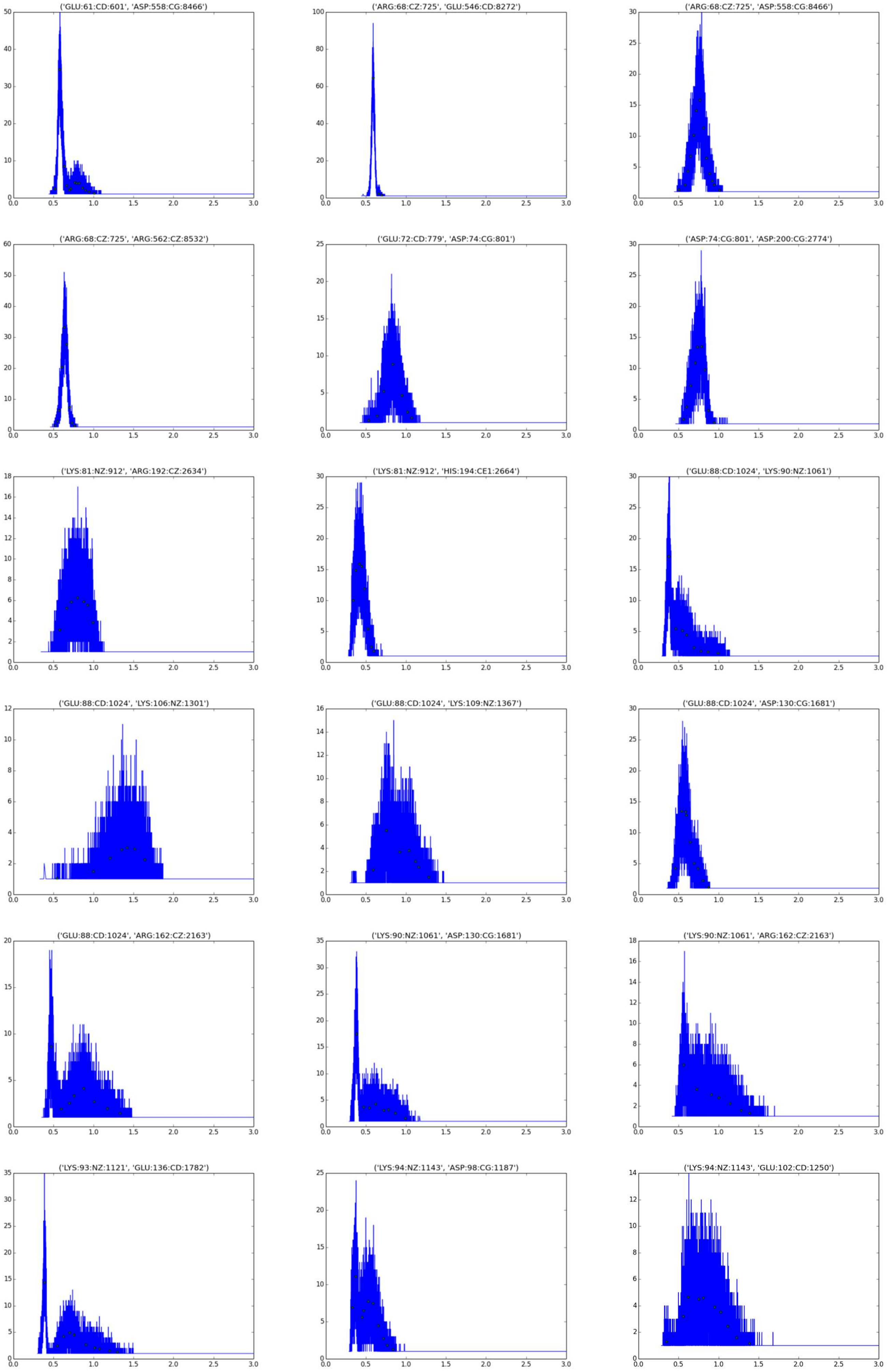
('GLU:61:CD:601', 'ASP:558:CG:8466')
('ARG:68:CZ:725', 'GLU:546:CD:8272')
('ARG:68:CZ:725', 'ASP:558:CG:8466')
('ARG:68:CZ:725', 'ARG:562:CZ:8532')
('GLU:72:CD:779', 'ASP:74:CG:801')
('ASP:74:CG:801', 'ASP:200:CG:2774')
('LYS:81:NZ:912', 'ARG:192:CZ:2634')
('LYS:81:NZ:912', 'HIS:194:CE1:2664')
('GLU:88:CD:1024', 'LYS:90:NZ:1061')
('GLU:88:CD:1024', 'LYS:106:NZ:1301')
('GLU:88:CD:1024', 'LYS:109:NZ:1367')
('GLU:88:CD:1024', 'ASP:130:CG:1681')
('GLU:88:CD:1024', 'ARG:162:CZ:2163')
('LYS:90:NZ:1061', 'ASP:130:CG:1681')
('LYS:90:NZ:1061', 'ARG:162:CZ:2163')
('LYS:93:NZ:1121', 'GLU:136:CD:1782')
('LYS:94:NZ:1143', 'ASP:98:CG:1187')
('LYS:94:NZ:1143', 'GLU:102:CD:1250')

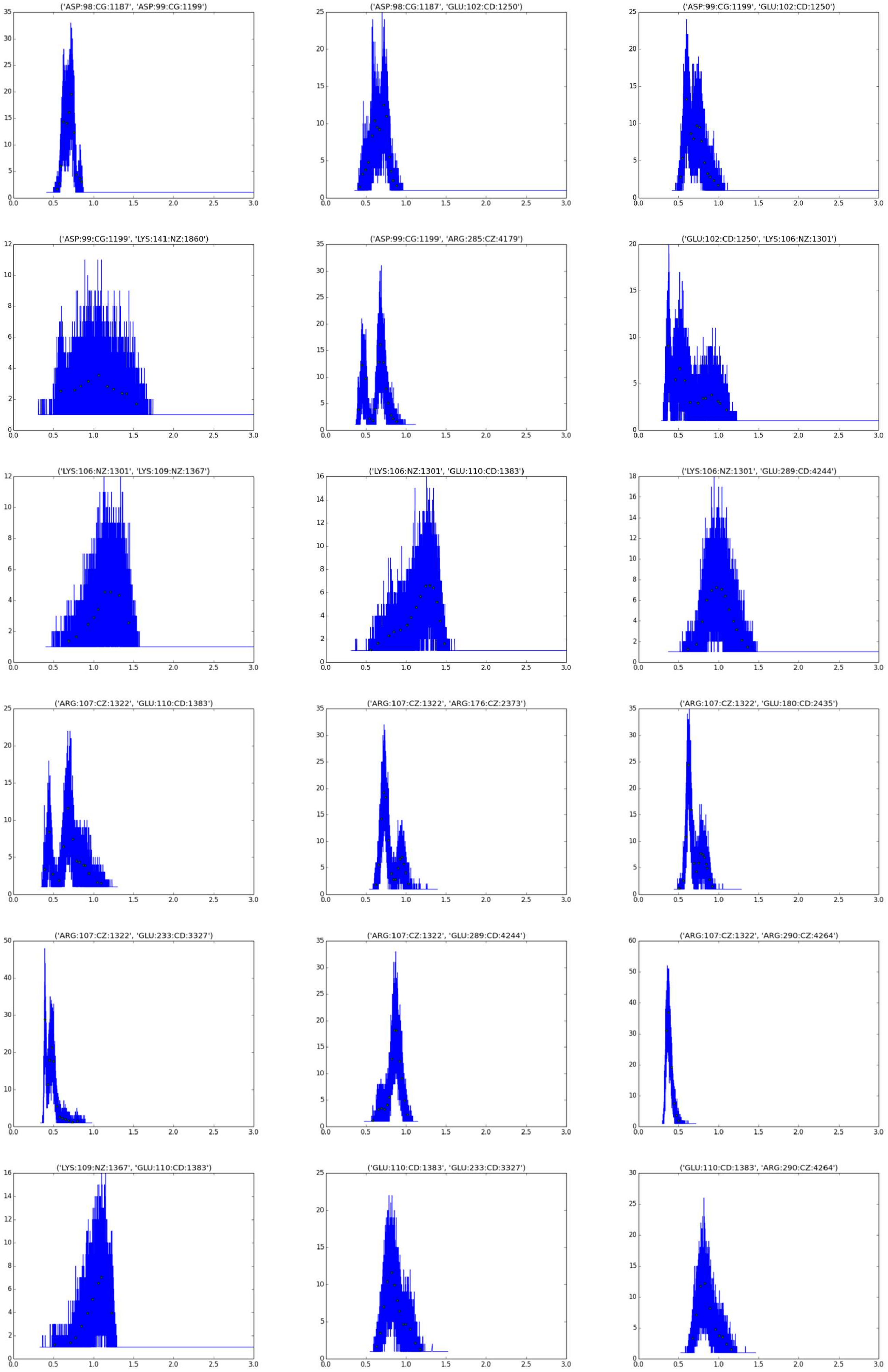
('ASP:98:CG:1187', 'ASP:99:CG:1199')
('ASP:98:CG:1187', 'GLU:102:CD:1250')
('ASP:99:CG:1199', 'GLU:102:CD:1250')
('ASP:99:CG:1199', 'LYS:141:NZ:1860')
('ASP:99:CG:1199', 'ARG:285:CZ:4179')
('GLU:102:CD:1250', 'LYS:106:NZ:1301')
('LYS:106:NZ:1301', 'LYS:109:NZ:1367')
('LYS:106:NZ:1301', 'GLU:110:CD:1383')
('LYS:106:NZ:1301', 'GLU:289:CD:4244')
('ARG:107:CZ:1322', 'GLU:110:CD:1383')
('ARG:107:CZ:1322', 'ARG:176:CZ:2373')
('ARG:107:CZ:1322', 'GLU:180:CD:2435')
('ARG:107:CZ:1322', 'GLU:233:CD:3327')
('ARG:107:CZ:1322', 'GLU:289:CD:4244')
('ARG:107:CZ:1322', 'ARG:290:CZ:4264')
('LYS:109:NZ:1367', 'GLU:110:CD:1383')
('GLU:110:CD:1383', 'GLU:233:CD:3327')
('GLU:110:CD:1383', 'ARG:290:CZ:4264')

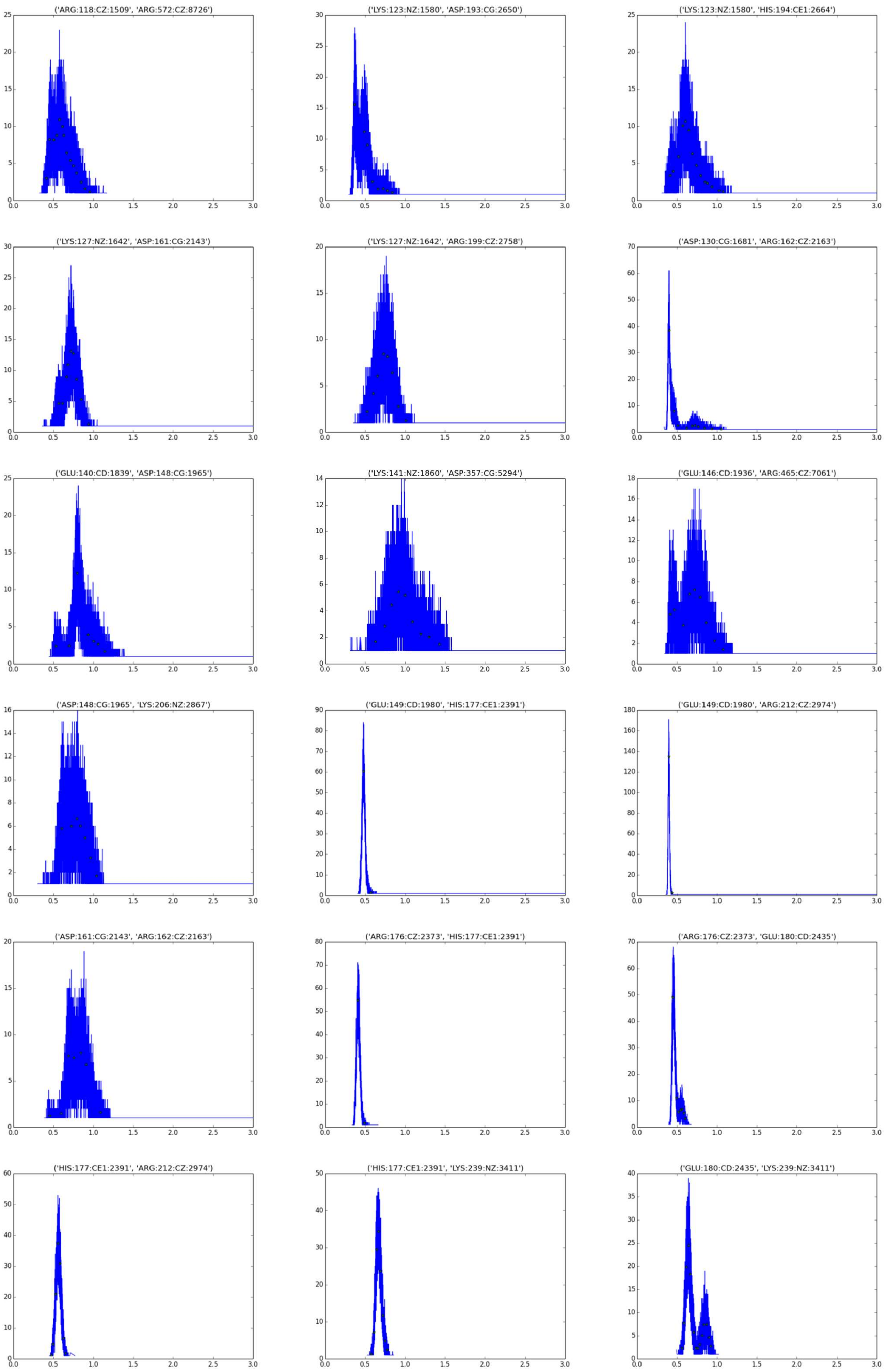
('ARG:118:CZ:1509', 'ARG:572:CZ:8726')
('LYS:123:NZ:1580', 'ASP:193:CG:2650')
('LYS:123:NZ:1580', 'HIS:194:CE1:2664')
('LYS:127:NZ:1642', 'ASP:161:CG:2143')
('LYS:127:NZ:1642', 'ARG:199:CZ:2758')
('ASP:130:CG:1681', 'ARG:162:CZ:2163')
('GLU:140:CD:1839', 'ASP:148:CG:1965')
('LYS:141:NZ:1860', 'ASP:357:CG:5294')
('GLU:146:CD:1936', 'ARG:465:CZ:7061')
('ASP:148:CG:1965', 'LYS:206:NZ:2867')
('GLU:149:CD:1980', 'HIS:177:CE1:2391')
('GLU:149:CD:1980', 'ARG:212:CZ:2974')
('ASP:161:CG:2143', 'ARG:162:CZ:2163')
('ARG:176:CZ:2373', 'HIS:177:CE1:2391')
('ARG:176:CZ:2373', 'GLU:180:CD:2435')
('HIS:177:CE1:2391', 'ARG:212:CZ:2974')
('HIS:177:CE1:2391', 'LYS:239:NZ:3411')
('GLU:180:CD:2435', 'LYS:239:NZ:3411')

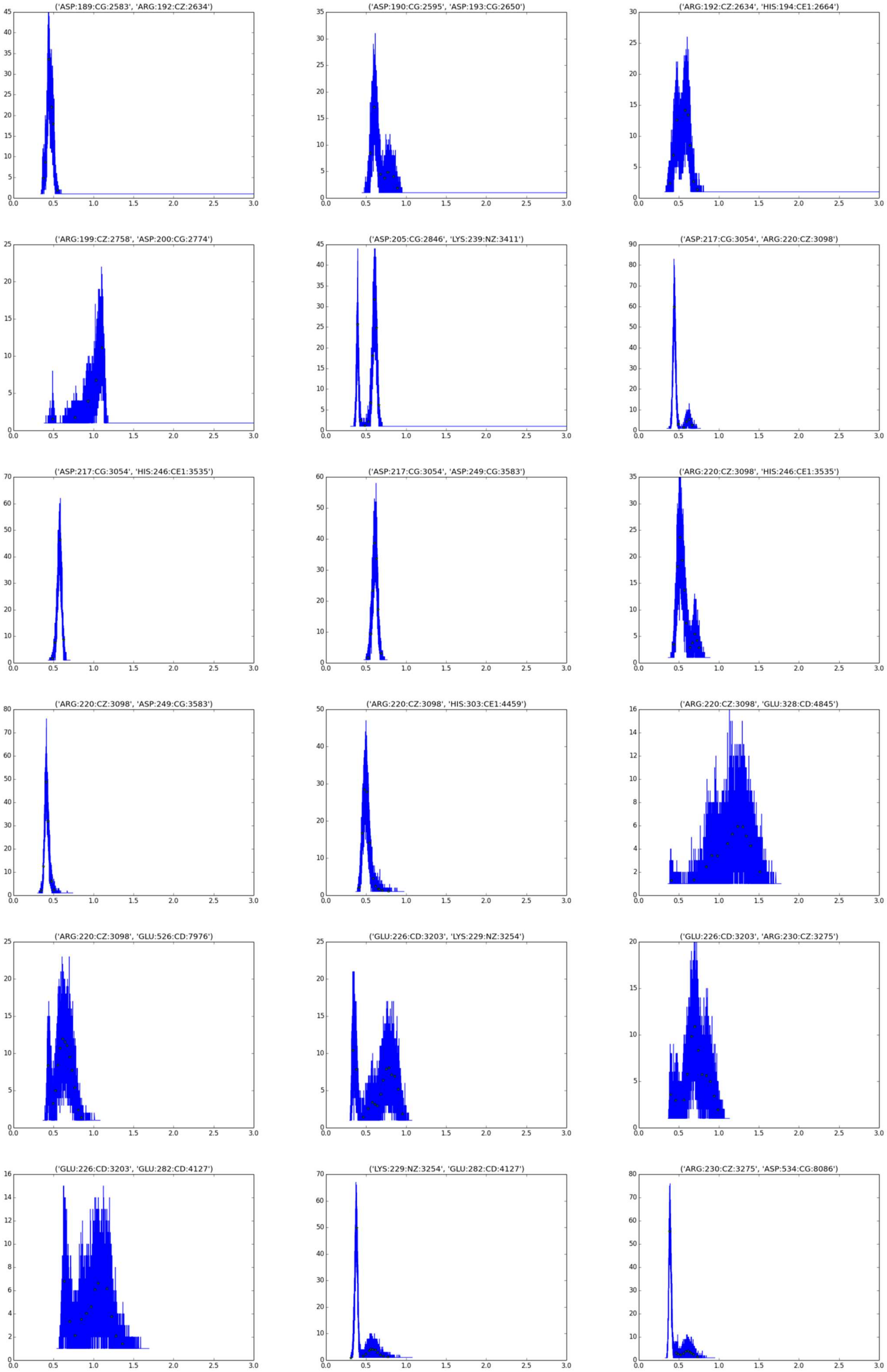
('ASP:189:CG:2583', 'ARG:192:CZ:2634')
('ASP:190:CG:2595', 'ASP:193:CG:2650')
('ARG:192:CZ:2634', 'HIS:194:CE1:2664')
('ARG:199:CZ:2758', 'ASP:200:CG:2774')
('ASP:205:CG:2846', 'LYS:239:NZ:3411')
('ASP:217:CG:3054', 'ARG:220:CZ:3098')
('ASP:217:CG:3054', 'HIS:246:CE1:3535')
('ASP:217:CG:3054', 'ASP:249:CG:3583')
('ARG:220:CZ:3098', 'HIS:246:CE1:3535')
('ARG:220:CZ:3098', 'ASP:249:CG:3583')
('ARG:220:CZ:3098', 'HIS:303:CE1:4459')
('ARG:220:CZ:3098', 'GLU:328:CD:4845')
('ARG:220:CZ:3098', 'GLU:526:CD:7976')
('GLU:226:CD:3203', 'LYS:229:NZ:3254')
('GLU:226:CD:3203', 'ARG:230:CZ:3275')
('GLU:226:CD:3203', 'GLU:282:CD:4127')
('LYS:229:NZ:3254', 'GLU:282:CD:4127')
('ARG:230:CZ:3275', 'ASP:534:CG:8086')

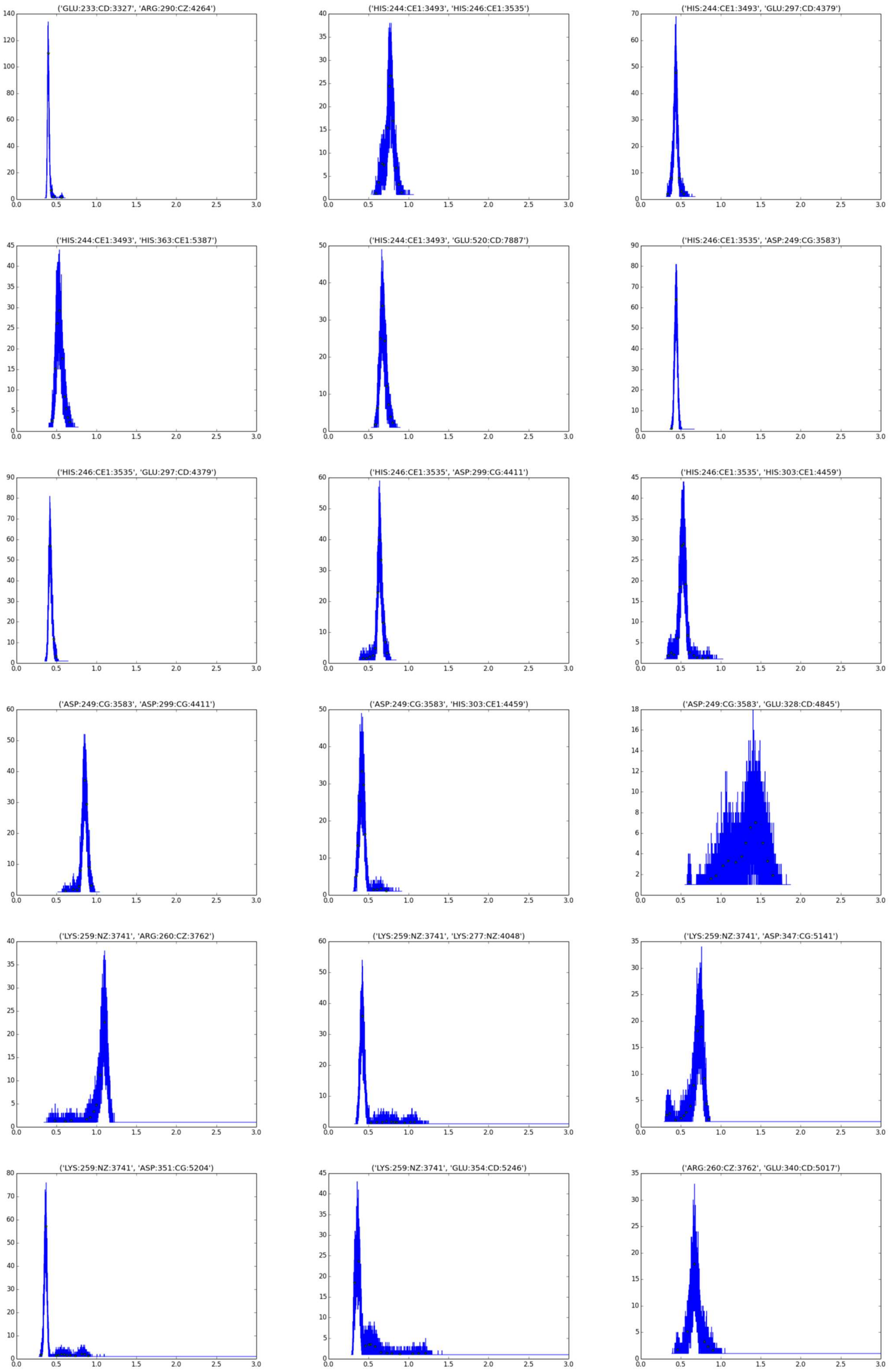
('GLU:233:CD:3327', 'ARG:290:CZ:4264')
('HIS:244:CE1:3493', 'HIS:246:CE1:3535')
('HIS:244:CE1:3493', 'GLU:297:CD:4379')
('HIS:244:CE1:3493', 'HIS:363:CE1:5387')
('HIS:244:CE1:3493', 'GLU:520:CD:7887')
('HIS:246:CE1:3535', 'ASP:249:CG:3583')
('HIS:246:CE1:3535', 'GLU:297:CD:4379')
('HIS:246:CE1:3535', 'ASP:299:CG:4411')
('HIS:246:CE1:3535', 'HIS:303:CE1:4459')
('ASP:249:CG:3583', 'ASP:299:CG:4411')
('ASP:249:CG:3583', 'HIS:303:CE1:4459')
('ASP:249:CG:3583', 'GLU:328:CD:4845')
('LYS:259:NZ:3741', 'ARG:260:CZ:3762')
('LYS:259:NZ:3741', 'LYS:277:NZ:4048')
('LYS:259:NZ:3741', 'ASP:347:CG:5141')
('LYS:259:NZ:3741', 'ASP:351:CG:5204')
('LYS:259:NZ:3741', 'GLU:354:CD:5246')
('ARG:260:CZ:3762', 'GLU:340:CD:5017')

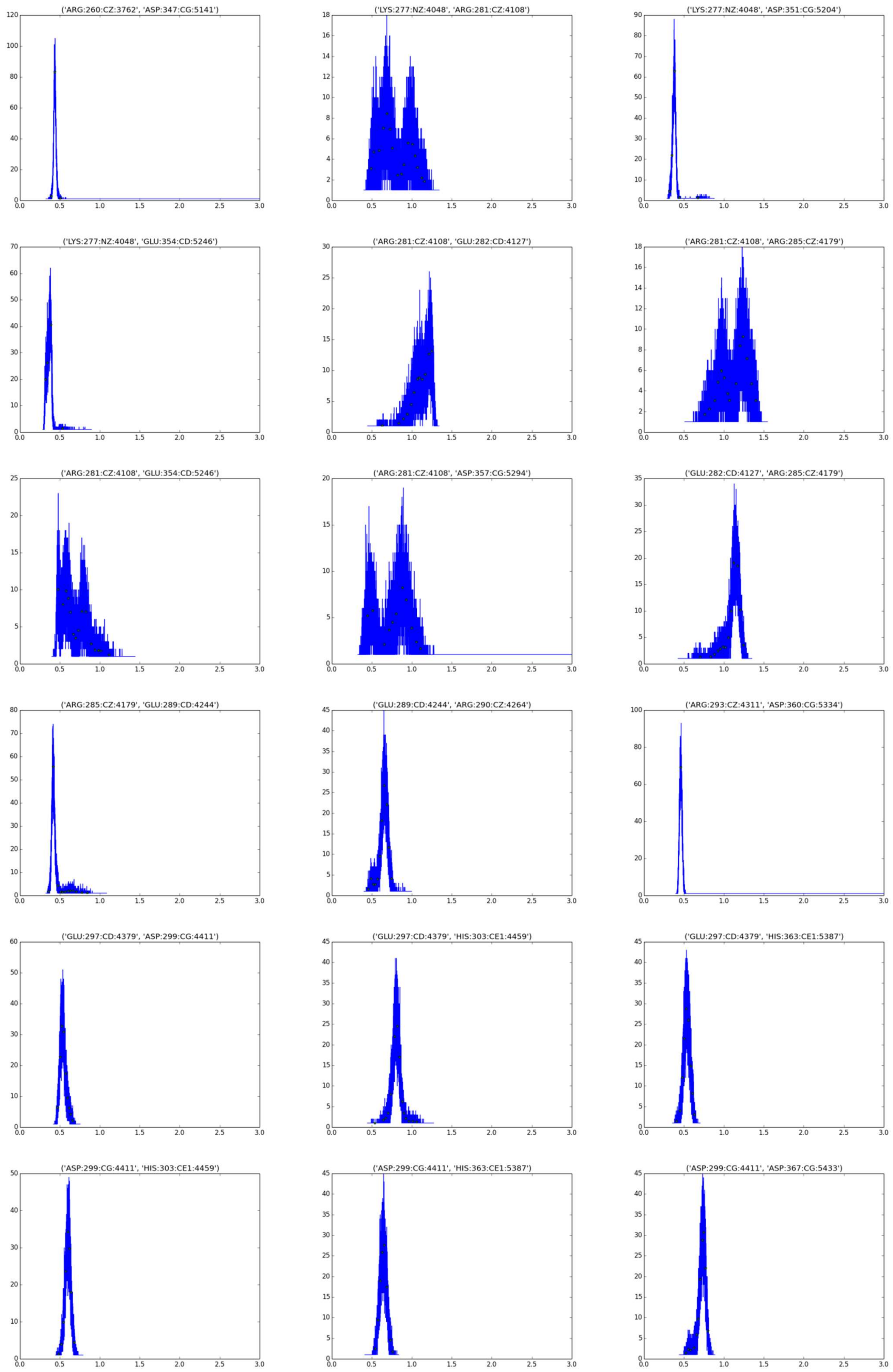
('ARG:260:CZ:3762', 'ASP:347:CG:5141')
('LYS:277:NZ:4048', 'ARG:281:CZ:4108')
('LYS:277:NZ:4048', 'ASP:351:CG:5204')
('LYS:277:NZ:4048', 'GLU:354:CD:5246')
('ARG:281:CZ:4108', 'GLU:282:CD:4127')
('ARG:281:CZ:4108', 'ARG:285:CZ:4179')
('ARG:281:CZ:4108', 'GLU:354:CD:5246')
('ARG:281:CZ:4108', 'ASP:357:CG:5294')
('GLU:282:CD:4127', 'ARG:285:CZ:4179')
('ARG:285:CZ:4179', 'GLU:289:CD:4244')
('GLU:289:CD:4244', 'ARG:290:CZ:4264')
('ARG:293:CZ:4311', 'ASP:360:CG:5334')
('GLU:297:CD:4379', 'ASP:299:CG:4411')
('GLU:297:CD:4379', 'HIS:303:CE1:4459')
('GLU:297:CD:4379', 'HIS:363:CE1:5387')
('ASP:299:CG:4411', 'HIS:303:CE1:4459')
('ASP:299:CG:4411', 'HIS:363:CE1:5387')
('ASP:299:CG:4411', 'ASP:367:CG:5433')

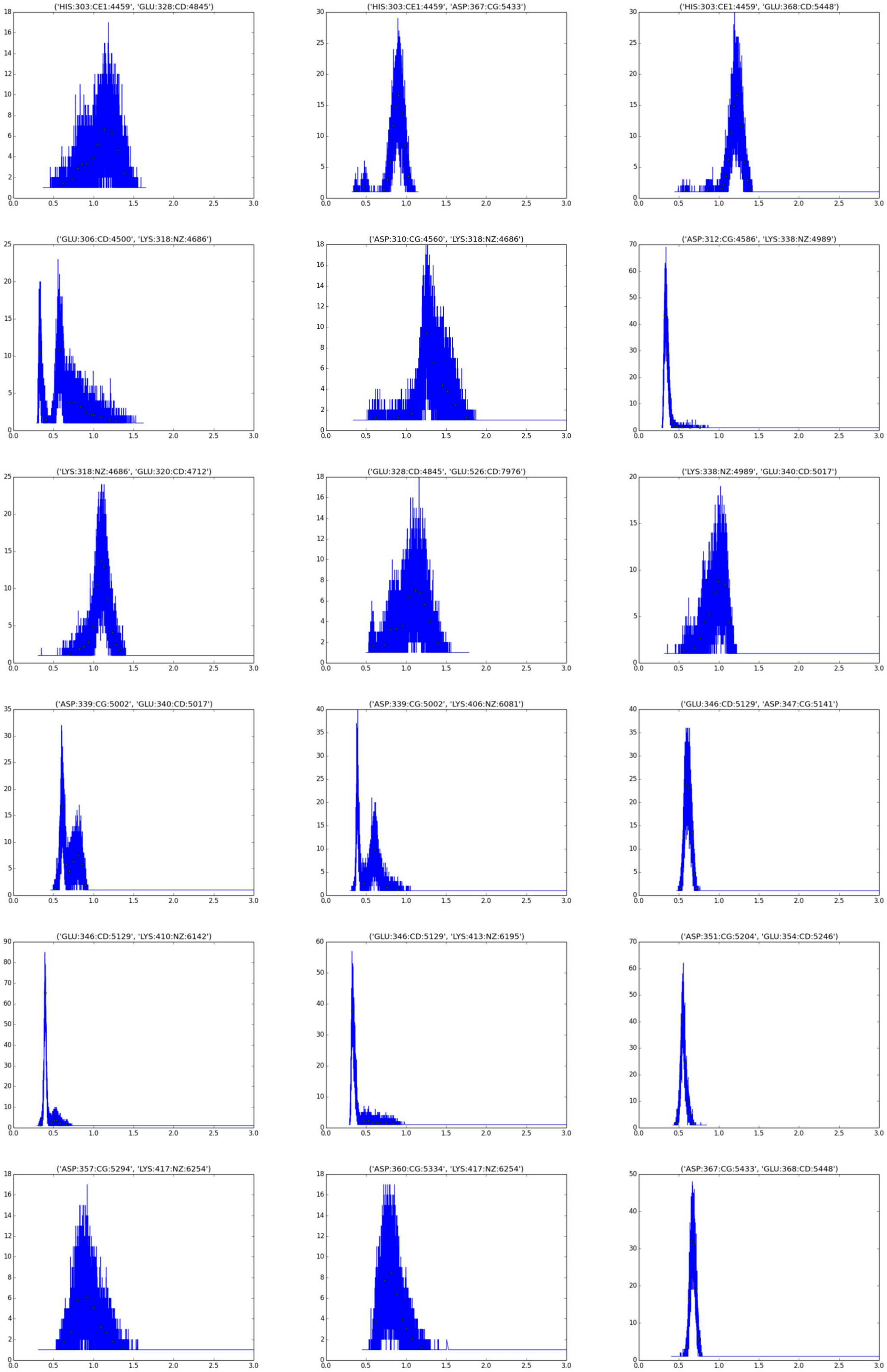
('HIS:303:CE1:4459', 'GLU:328:CD:4845')
('HIS:303:CE1:4459', 'ASP:367:CG:5433')
('HIS:303:CE1:4459', 'GLU:368:CD:5448')
('GLU:306:CD:4500', 'LYS:318:NZ:4686')
('ASP:310:CG:4560', 'LYS:318:NZ:4686')
('ASP:312:CG:4586', 'LYS:338:NZ:4989')
('LYS:318:NZ:4686', 'GLU:320:CD:4712')
('GLU:328:CD:4845', 'GLU:526:CD:7976')
('LYS:338:NZ:4989', 'GLU:340:CD:5017')
('ASP:339:CG:5002', 'GLU:340:CD:5017')
('ASP:339:CG:5002', 'LYS:406:NZ:6081')
('GLU:346:CD:5129', 'ASP:347:CG:5141')
('GLU:346:CD:5129', 'LYS:410:NZ:6142')
('GLU:346:CD:5129', 'LYS:413:NZ:6195')
('ASP:351:CG:5204', 'GLU:354:CD:5246')
('ASP:357:CG:5294', 'LYS:417:NZ:6254')
('ASP:360:CG:5334', 'LYS:417:NZ:6254')
('ASP:367:CG:5433', 'GLU:368:CD:5448')

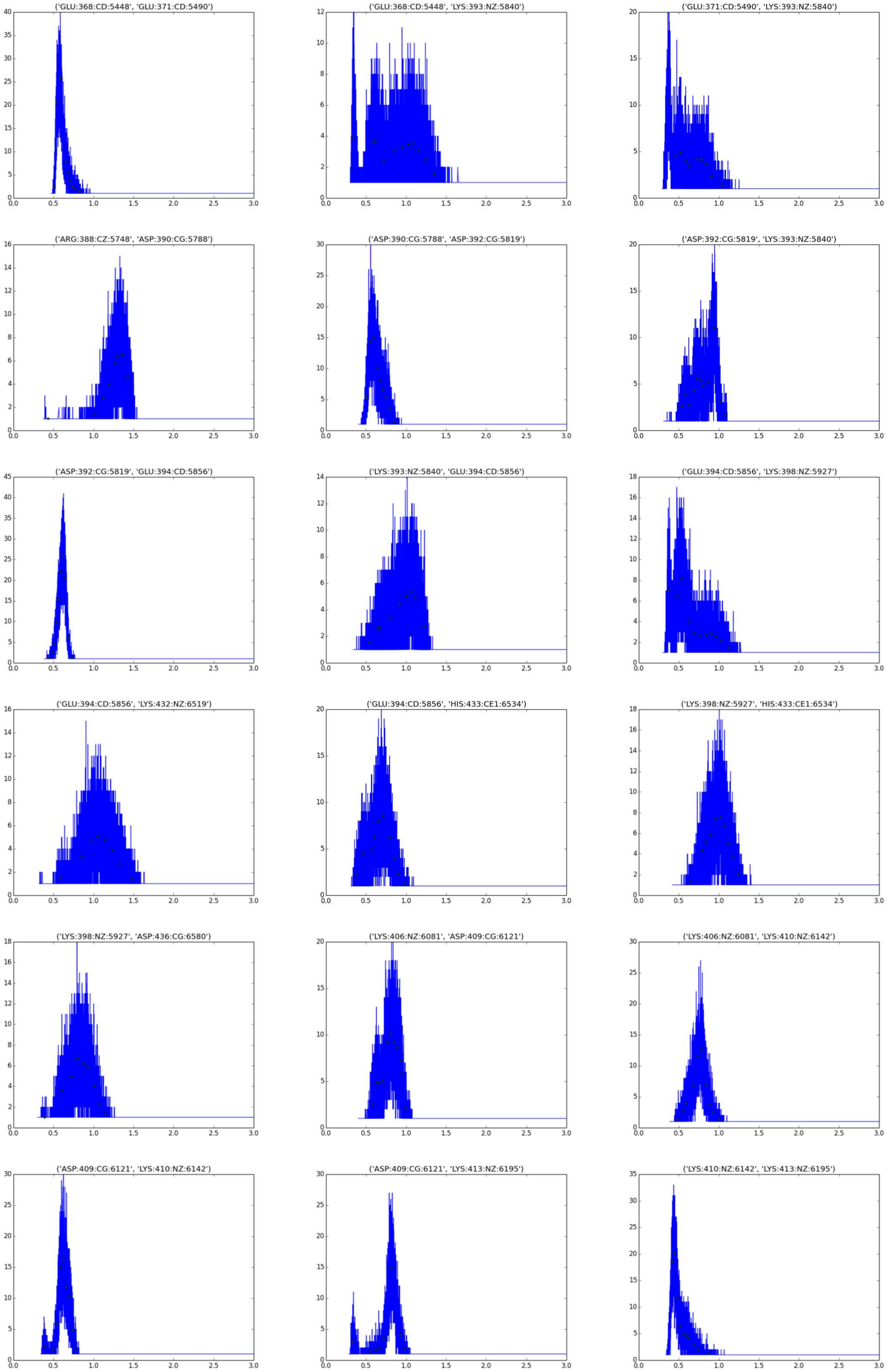
('GLU:368:CD:5448', 'GLU:371:CD:5490')
('GLU:368:CD:5448', 'LYS:393:NZ:5840')
('GLU:371:CD:5490', 'LYS:393:NZ:5840')
('ARG:388:CZ:5748', 'ASP:390:CG:5788')
('ASP:390:CG:5788', 'ASP:392:CG:5819')
('ASP:392:CG:5819', 'LYS:393:NZ:5840')
('ASP:392:CG:5819', 'GLU:394:CD:5856')
('LYS:393:NZ:5840', 'GLU:394:CD:5856')
('GLU:394:CD:5856', 'LYS:398:NZ:5927')
('GLU:394:CD:5856', 'LYS:432:NZ:6519')
('GLU:394:CD:5856', 'HIS:433:CE1:6534')
('LYS:398:NZ:5927', 'HIS:433:CE1:6534')
('LYS:398:NZ:5927', 'ASP:436:CG:6580')
('LYS:406:NZ:6081', 'ASP:409:CG:6121')
('LYS:406:NZ:6081', 'LYS:410:NZ:6142')
('ASP:409:CG:6121', 'LYS:410:NZ:6142')
('ASP:409:CG:6121', 'LYS:413:NZ:6195')
('LYS:410:NZ:6142', 'LYS:413:NZ:6195')

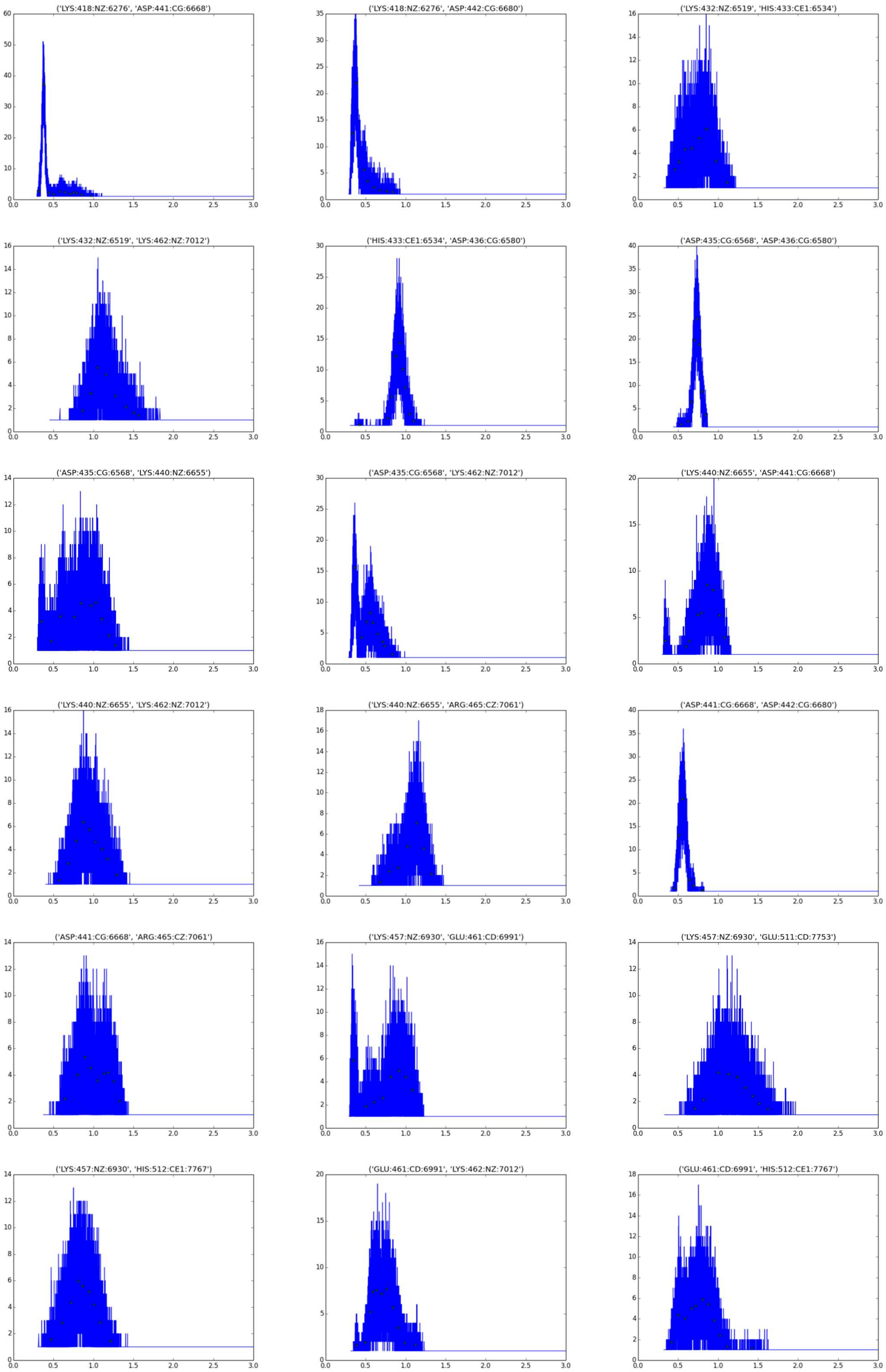
('LYS:418:NZ:6276', 'ASP:441:CG:6668')
('LYS:418:NZ:6276', 'ASP:442:CG:6680')
('LYS:432:NZ:6519', 'HIS:433:CE1:6534')
('LYS:432:NZ:6519', 'LYS:462:NZ:7012')
('HIS:433:CE1:6534', 'ASP:436:CG:6580')
('ASP:435:CG:6568', 'ASP:436:CG:6580')
('ASP:435:CG:6568', 'LYS:440:NZ:6655')
('ASP:435:CG:6568', 'LYS:462:NZ:7012')
('LYS:440:NZ:6655', 'ASP:441:CG:6668')
('LYS:440:NZ:6655', 'LYS:462:NZ:7012')
('LYS:440:NZ:6655', 'ARG:465:CZ:7061')
('ASP:441:CG:6668', 'ASP:442:CG:6680')
('ASP:441:CG:6668', 'ARG:465:CZ:7061')
('LYS:457:NZ:6930', 'GLU:461:CD:6991')
('LYS:457:NZ:6930', 'GLU:511:CD:7753')
('LYS:457:NZ:6930', 'HIS:512:CE1:7767')
('GLU:461:CD:6991', 'LYS:462:NZ:7012')
('GLU:461:CD:6991', 'HIS:512:CE1:7767')

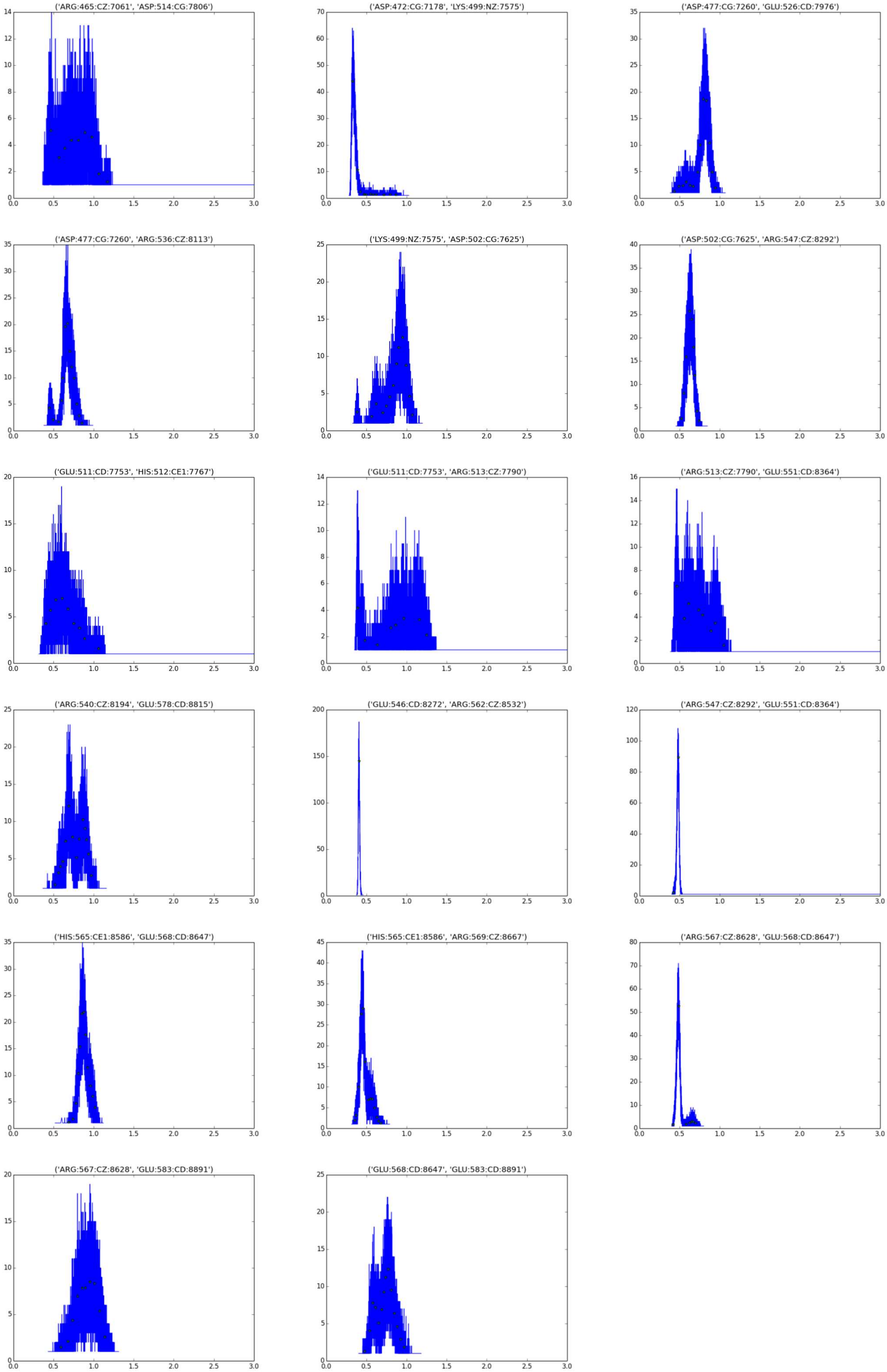
('ARG:465:CZ:7061', 'ASP:514:CG:7806')
('ASP:472:CG:7178', 'LYS:499:NZ:7575')
('ASP:477:CG:7260', 'GLU:526:CD:7976')
('ASP:477:CG:7260', 'ARG:536:CZ:8113')
('LYS:499:NZ:7575', 'ASP:502:CG:7625')
('ASP:502:CG:7625', 'ARG:547:CZ:8292')
('GLU:511:CD:7753', 'HIS:512:CE1:7767')
('GLU:511:CD:7753', 'ARG:513:CZ:7790')
('ARG:513:CZ:7790', 'GLU:551:CD:8364')
('ARG:540:CZ:8194', 'GLU:578:CD:8815')
('GLU:546:CD:8272', 'ARG:562:CZ:8532')
('ARG:547:CZ:8292', 'GLU:551:CD:8364')
('HIS:565:CE1:8586', 'GLU:568:CD:8647')
('HIS:565:CE1:8586', 'ARG:569:CZ:8667')
('ARG:567:CZ:8628', 'GLU:568:CD:8647')
('ARG:567:CZ:8628', 'GLU:583:CD:8891')
('GLU:568:CD:8647', 'GLU:583:CD:8891')

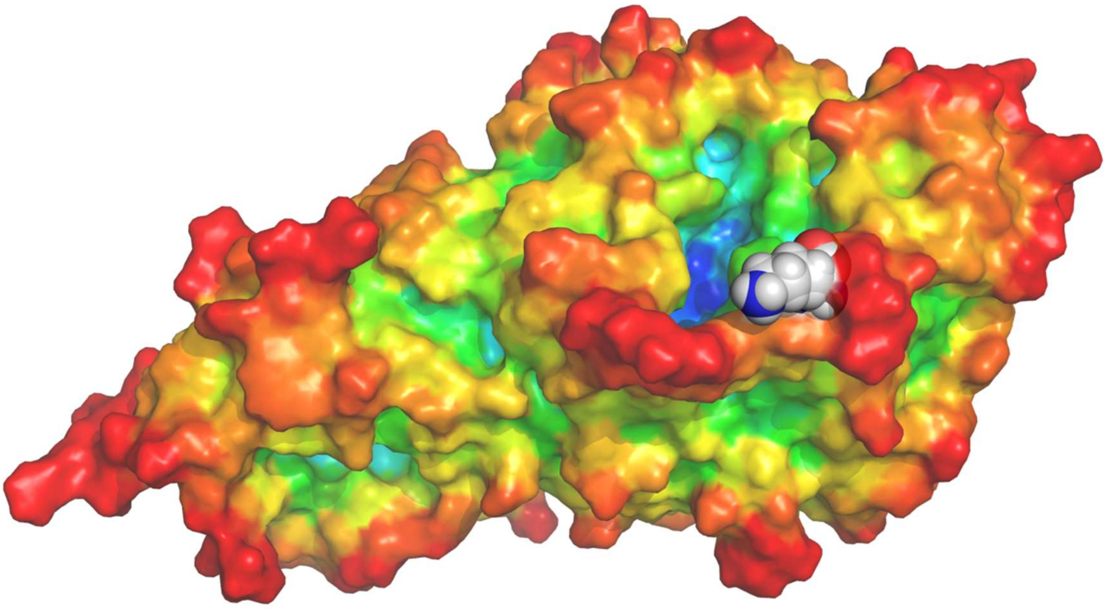

**Figure S6**: The structure of *Ostrinia furnacalis* chitinolytic enzyme: protein surface is colored according to 3D atom depth values, to highlight the catalytic site; K323 side chain is shown with space-fill representation.
